# The IRE1-bZIP60 branch of Unfolded Protein Response is required for *Arabidopsis* immune response to *Botrytis cinerea*

**DOI:** 10.1101/2023.10.18.562849

**Authors:** Blanchard Cécile, Aimé Sébastien, Ducloy Amélie, Hichami Siham, Azzopardi Marianne², Cacas Jean-Luc, Lamotte Olivier

## Abstract

The Unfolded Protein Response (UPR) is a retrograde signalling pathway which is activated when endoplasmic reticulum (ER) proteostasis is disturbed. Here, we have investigated by reverse genetics the contribution of such pathway in *Arabidopsis thaliana* response to two necrotrophic fungi of agricultural importance, *Botrytis cinere*a which is responsible for the development of grey mold disease, and *Alternaria brassicicola* which triggers black spot disease. We found that the branch of UPR dependent on the INOSITOL-REQUIRING ENZYME 1 (IRE1) and the transcription factor (TF) bZIP60 is required to restrict foliar necrotic symptoms induced by both fungi. Accordingly, focussing on *B. cinerea*, we provided evidence for the production of the active bZIP60 form during infection. This activation was accompanied by an increased expression of UPR-responsive genes coding for ER-localized chaperones and co-chaperones that belong to the ER-Quality Control (ER-QC) system. Furthermore, mutants deficient for two ER-QC components were also more susceptible to infection. By contrast, investigating the involvement of CELL DIVISION CYCLE 48 (CDC48) AAA+-ATPAses that assist ER-Associated Degradation (ERAD) pathway for disposal of luminal unfolded proteins, we showed that a series of mutants and transgenics are more resistant to grey mold disease. Seeking for molecular insights into how the ER could shape Arabidopsis immune response to *B. cinerea*, we quantified the expression of defence gene and cell death markers in single *bzip60* and double *ire1* mutants. However, none of those genes were mis-regulated in mutant genetic backgrounds, indicating that IRE1-bZIP60 branch of UPR modulates the Arabidopsis response to *B. cinerea* by a yet-to-be-identified mechanism. Interestingly, we identified the NAC053/NTL4 TF as a potential actor of this unknown mechanism, linking the UPR and proteasome stress regulon.

**Author summary:** Necrotrophic fungi are one of the most economically significant plant pathogens worldwide, inflicting massive pre- and post-harvest losses on a wide range of fruit and vegetable crops. They adopt a necrotrophic lifestyle, deriving their nutrients predominantly from dead plant tissues to complete their life cycle. *Botrytis cinerea* is the causal agent of grey mold and no plant shows complete resistance towards this pathogen. The use of genetic models such as the plant *Arabidopsis thaliana* has partially enabled the understanding of the immunity mechanisms involved in the plant’s response to *B. cinerea*. Our work provides new insights into the cellular mechanisms of how plants cope with this pathogen. In this context, by means of a reverse genetic approach, we explored the role of the Unfolded Protein Response (UPR), a cell signalling pathway regulating protein homeostasis within the endoplasmic reticulum (ER) and thus protecting cells from a harmful over-accumulation of aberrant or misfolded proteins.

## Introduction

The maintenance of protein homeostasis is one of the cornerstones of cellular functions. It involves the precise regulation of translation, the folding of newly synthesized proteins, their post-translational modifications, their sorting and trafficking within the cell, and their degradation (1). The endoplasmic reticulum (ER) plays crucial roles in these processes, supporting the synthesis of one third to a quarter of total proteins in eukaryotic cells (2). Environmental factors, such as pathogen infection, intensify the workload on the protein folding machinery. When the endoplasmic reticulum (ER) is unable to meet the cell’s demand for protein folding, unfolded or misfolded polypeptides accumulate within the ER lumen, leading to a proteotoxic stress known as ER stress. This situation actuates the Unfolded Protein Response (UPR), a retrograde, ER-to-nucleus, signalling pathway, that is conserved across kingdoms. In Arabidopsis, the canonical UPR invokes three bZIP transcription factors (TF) that are anchored in ER membranes (bZIP17, bZIP28 and bZIP60) and made soluble, therefore active, through two distinct mechanisms when ER stress occurs. One mechanism is the non-conventional splicing of *bZIP60* mRNAs that results in a shift in the open reading frame, eventually causing the translated proteins to be devoid of their transmembrane domains. Such splicing process relies on the ribonuclease activity of two ER-resident transmembrane proteins, the INOSITOL-REQUIRING ENZYME 1A (IRE1A) and B (IRE1B) (3,4). When ER stress is persistent, IRE1 ribonuclease activity can also be employed to dispose of ER-associated mRNAs, a phenomenon known as Regulated IRE1-Dependant Decay (RIDD; (5,6); the latter of which reduces the amount of nascent polypeptides crossing ER membranes, thus preventing the accumulation of unfolded proteins within the ER lumen. The other activation mechanism of UPR involves the translocation of bZIP17 and/or bZIP28 TF to the Golgi apparatus through vesicular trafficking, their intramembrane proteolytic cleavage and subsequent release of their transcription-activating domains that can then reach the nucleus (7,8). Target genes of UPR TF encompasses those coding for chaperones, co-chaperones and additional factors involved in the ER-Quality Control (ER-QC) system, increasing ER folding capacity. This includes calreticulin (CRT) and calnexin (CNX), as well as BINDING-IMMUNOGLOBULIN PROTEINS (BiP) and its interacting proteins STROMAL CELL-DERIVED FACTOR 2 (SDF2) and ENDOPLASMIC RETICULUM DNAJ 3 (ERdj3) (9). In support of the transcriptional reprogramming mediated by the UPR TF, mis-folded and aberrant proteins can also be retro-translocated from ER lumen to cytoplasm where they are ubiquitinated for proteasomal degradation. This process, so-called ER-associated degradation (ERAD), involves the HRD1 (3-Hydroxy-3-methylglutaryl Reductase Degradation 1) complex and the AAA+-ATPase family of proteins CDC48 (Cell Division Control protein 48, (9). It aims at restoring ER proteostasis. When adaptive UPR pathway fails to resolve ER stress, a genetic program that ends with the death of malfunctioning cells is engaged. In this context, it has been demonstrated that ER stress-induced cell death (ERSID) is under the control of NAC089 TF (10) and requires several cathepsin B proteases to be executed (11).

A plethora of environmental situations can activate the UPR in plants, including heat, salt and drought stress, as well as pathogen infection (12,13). However, only a few data report on UPR involvement in the host plant response to necrotrophic pathogens. For instance, it was demonstrated that inoculating *Nicotiana attenuata* with *Alternaria alternata* resulted in the activation of UPR, and silencing *NaIRE1* or *NabZIP60* genes rendered plants more susceptible to the fungus (14). It was also showed that impairment of the IRE1/bZIP60 branch enhanced Arabidopsis susceptibility to *Drechslera gigantea*, another necrotrophic fungus (15). Although diseases caused by the necrotrophic fungi *Botrytis cinerea* and *Alternaria brassicicola* rank among the most devastating plant diseases worldwide, there are no data documenting the role of UPR in response to these two necrotrophic pathogens. *Botrytis cinerea* causes grey mold disease in a wide range of crops, and *Alternaria brassicicola* is responsible for black spot disease in numerous Brassica species. Both pathogens have the ability to infect the plant model *Arabidopsis thaliana*, making it a valuable tool for unravelling the intricate signalling pathways involved in plant immune response to these pathogens. Innate immunity in plants is orchestrated through signalling cascades involving key hormones like salicylic acid (SA), jasmonic acid (JA), and ethylene (ET), inducing significant modifications in gene expression that underlie enhanced resistance (16). The JA signalling pathway is commonly linked to the establishment of plant defence against necrotrophic fungus (17). Arabidopsis mutants with impaired JA perception, production or signalling are notably vulnerable to *B. cinerea* or *A. brassisicola* (18,19). SA is recognized as a pivotal hormone that initiates immune responses against biotrophic pathogens (16), but SA-mediated signalling has also been shown to be involved in resistance against *B. cinerea* in Arabidopsis (19) or against *Alternaria solani* in potato (20). For a comprehensive probing of hormonal signalling cascades, studying the expression profile of marker genes is a widely established method. Activation of JA-dependent signalling pathway is frequently assessed by investigating the expression of genes such as *PLANT DEFENSIN 1.2* (*PDF1.2*) or *OCTADECANOID-RESPONSIVE ARABIDOPSIS 59* (*ORA59*), whereas *PATHOGENESIS-RELATED-1 (PR1)* serves as a marker gene for SA (21). Another typical response of plants facing necrotrophic pathogens is the production in Arabidopsis of the phytoalexin camalexin through SA or JA signalling pathways, depending on the invading pathogen (22). The last step of camalexin synthesis is catalysed by the cytochrome P450 CYP71B15 (PHYTOALEXIN DEFICIENT 3, PAD3) which mRNA accumulation correlates with camalexin production (23). Camalexin has been shown to be toxic for *B. cinerea* and *A. brassicicola* (24,25), and *pad3* mutant shows enhanced susceptibility to both pathogens (19,26). During infection process, necrotrophic pathogens secrete compounds that enable rapid host cell death and disease spreading. *B. cinerea* produces cell-death inducing proteins, but also manipulates the plant regulated cell death to promote host cell death (27,28).

In this study, we explored how the ER could shape *Arabidopsis thaliana* immune response to necrotrophic pathogens, with a specific emphasis on *Botrytis cinerea*. The role of UPR, ER-QC and ERAD in this context was investigated using a reverse genetic approach in order to gain insights into molecular events that govern plant susceptibility and defence

## Results

### The IRE1-bZIP60 branch of Arabidopsis UPR restricts lesions induced by necrotrophic pathogens

To investigate a potential role of the UPR pathway in Arabidopsis response to necrotrophic fungi, we inoculated mutants defective in canonical UPR actors (*bzip17-1*, *bzip28-2*, *bzip60-3*, *ire1a-2 ire1b-4*) with *B. cinerea* or *A. brassicicola.* Mutant susceptibility to *B. cinerea* was examined 3 days upon drop-inoculation of leaves with a solution of conidia using the lesion diameter as proxy. The necrotic lesion diameter was averaging 5 mm in the WT genetic background, as previously reported (29), so was it in the *bzip28-2* mutant. By contrast, a significant decrease of 18 % in lesion size was recorded in *bzip17-1* mutant when compared to WT. Interestingly, both the single *bzip60-3* and double *ire1a-2 ire1b-4* mutants were more susceptible to the infection, displaying a 17 % increase in lesion diameter with respect to the WT genotype (Figure 1A, Supplemental Figure S1A). *A. brassicicola* susceptibility was also determined by inoculating leaves with a spore solution and lesion diameter were measured at 5 days post-inoculation. In WT plants, *A. brassicicola* induced necrotic lesions of 2.5 mm diameter in average. Both *bzip17-1* and *bzip28-2* mutations had no impact on symptoms. Genetic inactivation of *bZIP60* or *IRE1A* and *IRE1B* led to a 37 % or 27 % increase in lesion diameter with regard to WT, respectively (Figure 1B, Supplemental Figure S1B). Altogether, our data suggest that the IRE1-bZIP60 branch of UPR could be activated and participate either in defence mounting or PCD inhibition triggered by the two necrotrophic fungal pathogens tested.

**Figure 1:**
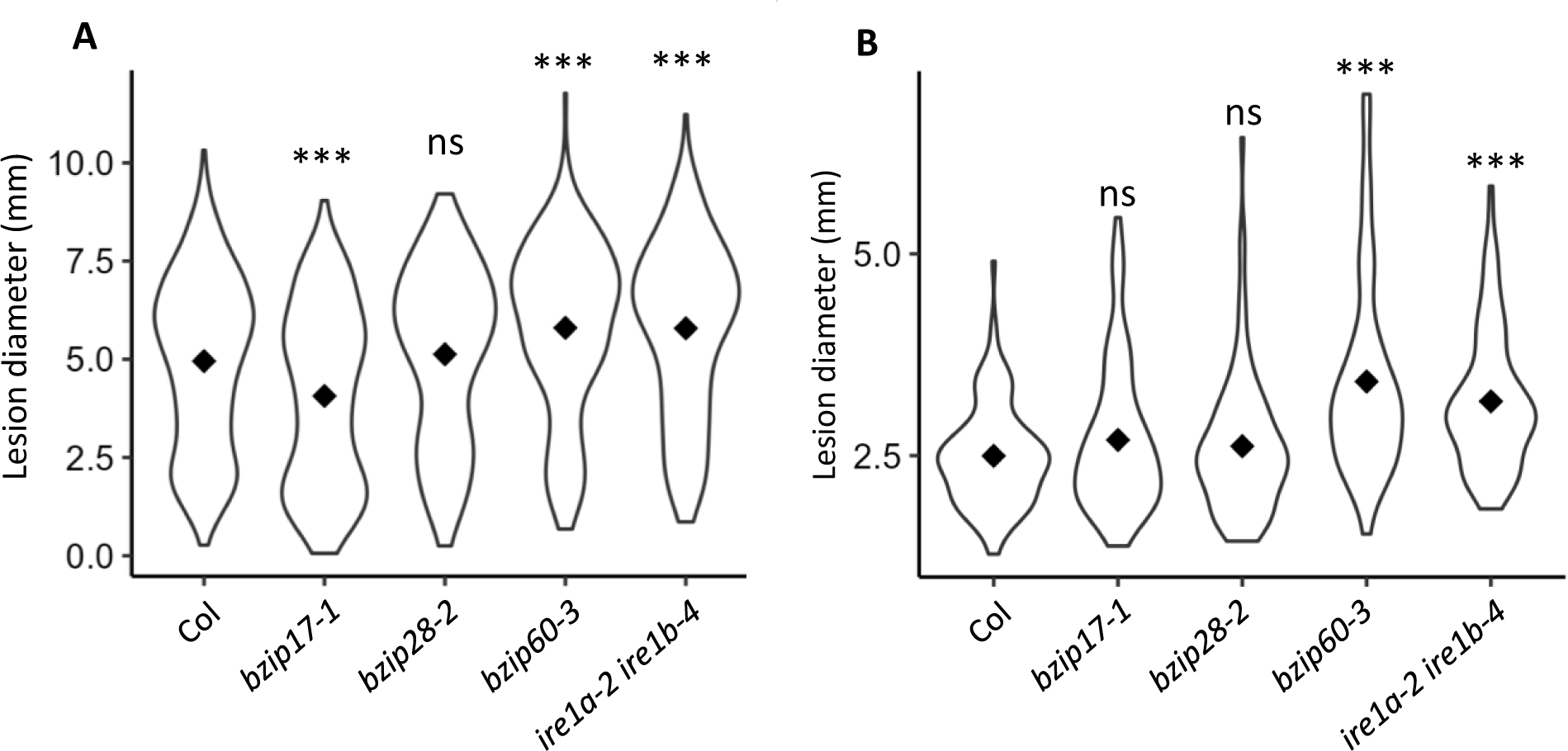
Mutants deficient for IRE1-bZIP60 branch of UPR show enhanced necrotic symptoms induced by *B. cinerea a*nd *A. brassicicola.* (A) Lesion diameters were measured 3 days post-inoculation with *B. cinerea* and (B) 5 days post- inoculation with *A. brassicicola.* Experiments were repeated seven to eight times for *B. cinerea* infection and four times for *A. brassicicola* infection depending on the genotype tested. Seven plants per genotype were infected in each experiment. Violin plots display the distribution of lesion diameters for all experiments and diamonds indicate the median. Stars indicate significant differences compared to the wild type and were assessed using Kruskal-Wallis’ method followed by Dunnett’s post-hoc test (*** p < 0.005; ns: no significant difference).

### The IRE1-bZIP60 branch of UPR is activated in response to *Botrytis cinerea*

To get into molecular mechanisms involving UPR, we focussed on the interaction between *A. thaliana* and *B. cinerea*. The expression of *IRE1A, IRE1B, bZIP17, bZIP28* and *bZIP60* was followed by RT-qPCR over time upon infection. For that purpose, a solution of *B. cinerea* spores prepared in PDB medium or PDB alone (mock) was sprayed on leaves and samples were collected at 16, 24, 48 and 72 h post-infection (hpi) for further processing. Those experimental conditions were validated by checking the expression of defence genes previously reported to be markedly induced in response to *B. cinerea*, *i.e. PAD3*, *PR1*, *PDF1.2a* and *PATATIN-LIKE PROTEIN 2 (PLP2)* (19,24,30). As expected, all four genes were up-regulated during infection (Supplemental Figure S2). As for UPR actors-encoded genes, *IRE*1A and *IRE1B* were not differentially expressed between mock and challenged conditions. A slight, but significant accumulation of *bZIP17* and *bZIP28* transcripts was only observed at 48 hpi (Figure 2). However, in accordance to mutant over-susceptibility to *B. cinerea* (Figure 1A), the strongest transcriptional response was observed for *bZIP60* gene with the unspliced mRNAs accumulating at early time points (16 and 24 hpi) whereas the spliced mRNA levels kept increasing from 16 hpi to 72 hpi. In addition to confirming the activation of the IRE1-bZIP60 branch, these results suggest that *B. cinerea* infection induces an ER stress.

**Figure 2:**
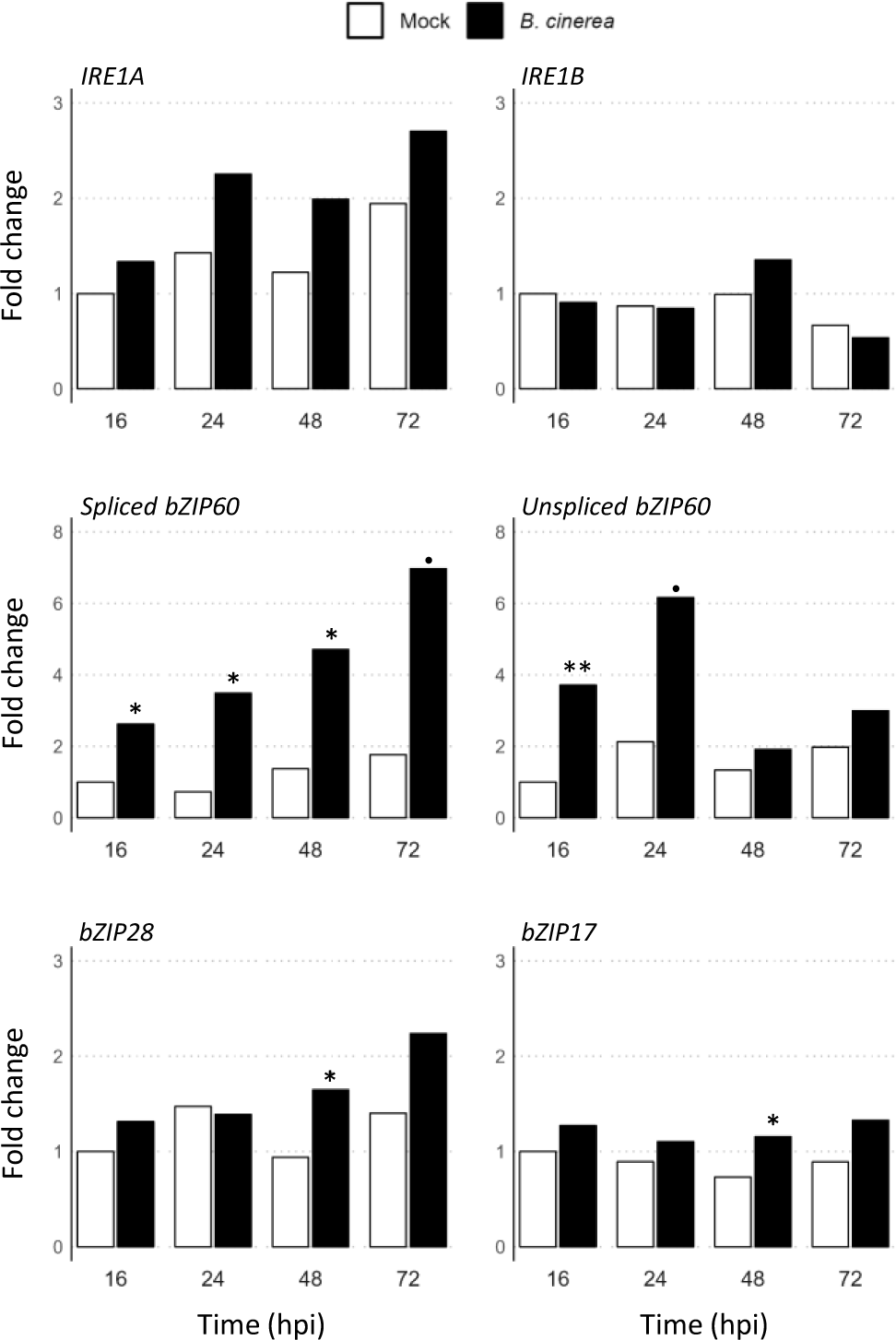
Expression kinetic of UPR genes in response to *B. cinerea*. Plants were sprayed with ¼ PDB solution (Mock, white bars) or with ¼ PDB solution containing *B. cinerea* spores (black bars). For each experiment, four leaves of three plants were collected at 16, 24, 48 and 72 hours post-inoculation. Level of transcripts coding IRE1A, IRE1B, bZIP17, bZIP28 and bZIP60 (unspliced or spliced forms) were quantified by RT-qPCR and normalized to those of two reference genes *AT4G26410* and *AT3G01150* (54). Experiments were repeated five times and mean expression for all experiments are presented. Significant differences between non-infected and infected plants were determined by a one-way ANOVA followed by a Tukey HSD Test (*** p<0.005; ** p<0.01; * p<0.05; p<0.1).

### Default in ER-QC machinery leads to enhanced necrotic lesions caused by *Botrytis cinerea*

We next determined by RT-qPCR the expression kinetic of ER stress gene hallmarks (*BIP1*, *BiP2*, *BiP3*, *ERDJ3A*, *ERDJ3B* and *SDF2*) in mock and inoculated WT plants (Figure 3A). Consistent with a proteotoxic ER stress induced by *B. cinerea*, all genes coding for ER-QC components were up-regulated upon infection, yet with distinct expression profiles. While *BIP1* and *BIP2* mRNAs were over-accumulating as early as 16 hpi, the expression of *BIP3* gene tends to increase at 48 hpi. The expression level of the three BiP-encoding genes were back to that of mock conditions at 72hpi. When compared to mock plants, the infection by *B. cinerea* was also stimulating the expression of *ERDJ3 A*, *ERDJ3B* and *SDF2* genes at 24 and/or 48 hpi depending on the gene tested. These results prompted us to investigate whether SDF2 and ERDJ3B could be involved in Arabidopsis response to *B. cinerea*. For that purpose, we used *erdj3b-1* and *sdf2-2* mutants (31) that we challenged with a spore solution and lesion diameters were measured at 3 days post-inoculation (Figure 3B, Supplemental Figure S3). Significant increases in lesion diameter were recorded for both mutants (24 % for *erdj3b-1* and 18 % for *sdf2-2*) in comparison to WT plants, indicating that they are more susceptible to *B. cinerea*. These data indicate that both SDF2 and ERDJ3B could be acting in the same response pathway as that of IRE1-bZIP60.

**Figure 3:**
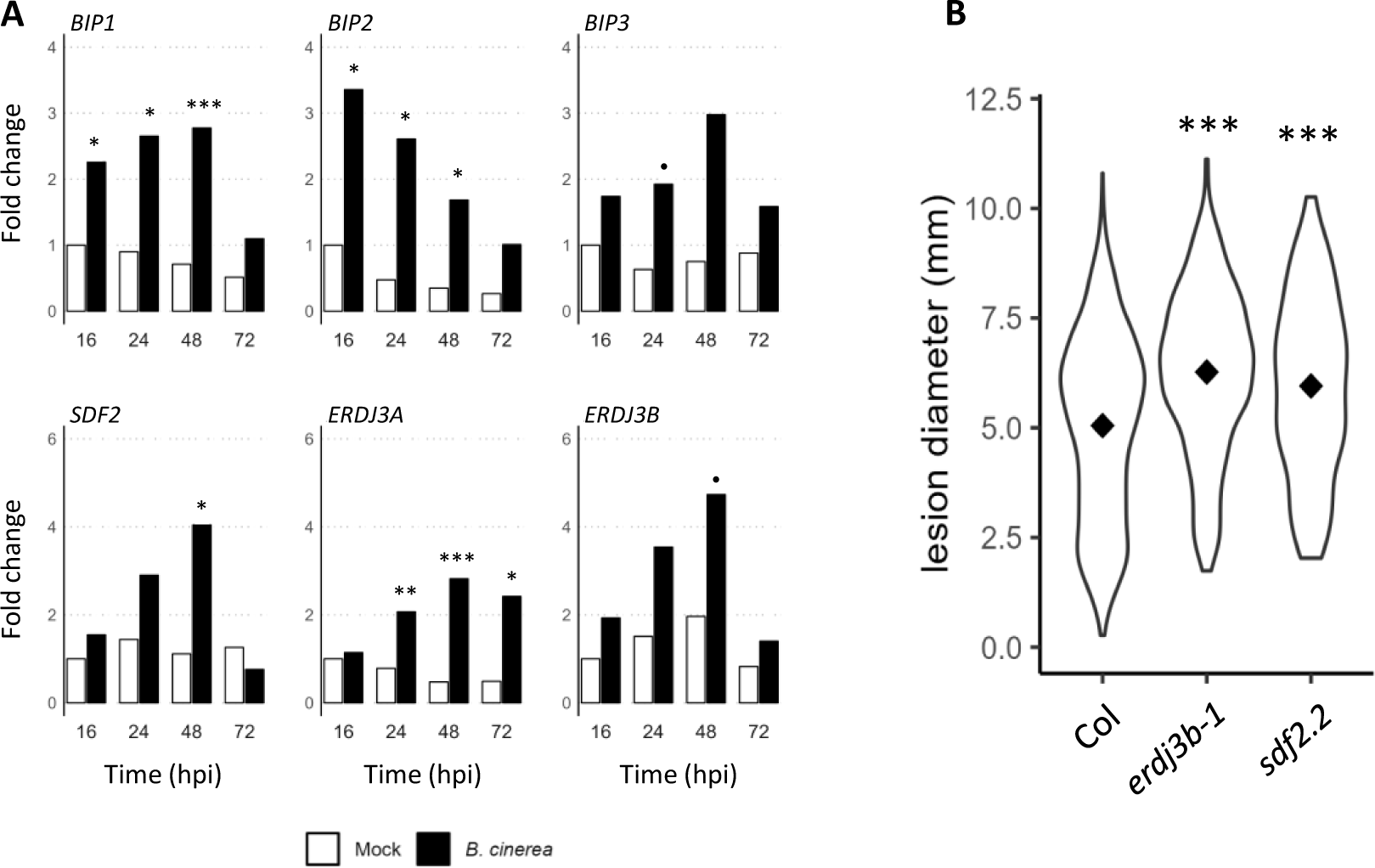
Defect in ER-QC machinery enhances *B. cinerea-*induced symptoms. (A) Expression of genes coding ER-QC machinery over *B. cinerea* infection. Plants were sprayed with ¼ PDB solution (Mock, white bars) or with ¼ PDB solution containing *B. cinerea* spores (black bars). For each experiment, four leaves of three plants were collected at 16, 24, 48 and 72 hours post-inoculation and used for total RNAs extraction. Transcript levels were quantified by RT-qPCR and normalized to those of two reference genes, *AT4G26410* and *AT3G01150* (54). The represented fold change is the mean of six independent experiments. Significant differences between non-infected and infected plants were determined by one-way ANOVA followed by a Tukey HSD post-hoc test (*** p<0.005; ** p<0.01; * p<0.05; p<0.1). (B) Disease phenotype of *erdj3b-1* and *sdf2-2* mutants infected by *B. cinerea*. Lesion diameters observed in wild-type (Col-0) and mutants were measured 3 days after *B. cinerea* infection. Experiments were repeated four times. Seven plants per genotype were infected in each experiment. Violin plots display the distribution of lesion diameters for all experiments and diamonds indicate the median. Stars indicate significant differences compared to the wild type and were using Kruskal-Wallis’ method followed by Dunnett’s post-hoc test (*** p < 0.005).

### Mutation in ERAD-related *CDC48* genes decreases susceptibility to *Botrytis cinerea*

The ERAD utilizes proteasome to dispose of misfolded proteins which have been retro-translocated from ER lumen to cytoplasm. With ER-QC system which is reinforced to increase ER folding capacity under ER stress, ERAD is part of the pro-adaptive mode of UPR that aims at restoring ER homeostasis. The CDC48 family of AAA+-ATPase proteins have been described as ERAD actors in yeast (32), mammals (33) and plants (34). To investigate a potential involvement of ERAD in response to *B. cinerea*, we first functionally characterized Arabidopsis CDC48 homologs. In Arabidopsis, CD48 proteins are, indeed, likely to be encoded by five genes (35). CDC48A, B and C are the closest sequence orthologs of human and yeast proteins, with which they share approximately 90 % similarity. By contrast, CDC48D and E are more divergent, exhibiting no or little conservation within their N and C-terminal domains (35,36). In accordance with its role in ERAD, CDC48A was shown to control the turnover of two immune receptors (36,37), as well as that of a mutated carboxypeptidase Y protein which is an ERAD substrate (34). It was also capable of complementing yeast *cdc48* conditional mutant (37). The four Arabidopsis homologs of *CDC48A* were thus tested by yeast functional complementation using the *Saccharomyces cerevisiae cdc48* mutant strains KFY189 and DBY2030. KFY189 is a temperature-sensitive mutant which does not grow at 37°C whereas DBY2030 is cold-sensitive and fail to grow at 16°C (38,39). Both strains were transformed with the yeast expression vector pDRF1-GW containing either the cDNA coding for CDC48A, B, C, D or E or with the empty vector alone. All transformed strains were able to grow under permissive conditions at 30°C (Figure 4A, C). As previously described (38), CDC48A complemented both strains under non-permissive conditions while the empty vector did not, validating our experimental setup (Figure 4B, D). Whereas *CDC48B* or *CDC48C* were able to restore the growth of DBY2030 strain at 16°C and that of KFY189 strain at 37°C (Figure 4B, D), *CDC48D* and *CDC48E* could not rescue the temperature-dependent phenotypes of both strains (Figure 4B, D). These data demonstrate that only CDC48A, B and C can functionally substitute to the yeast protein, and suggest a role for CDC48B and C in ERAD.

**Figure 4:**
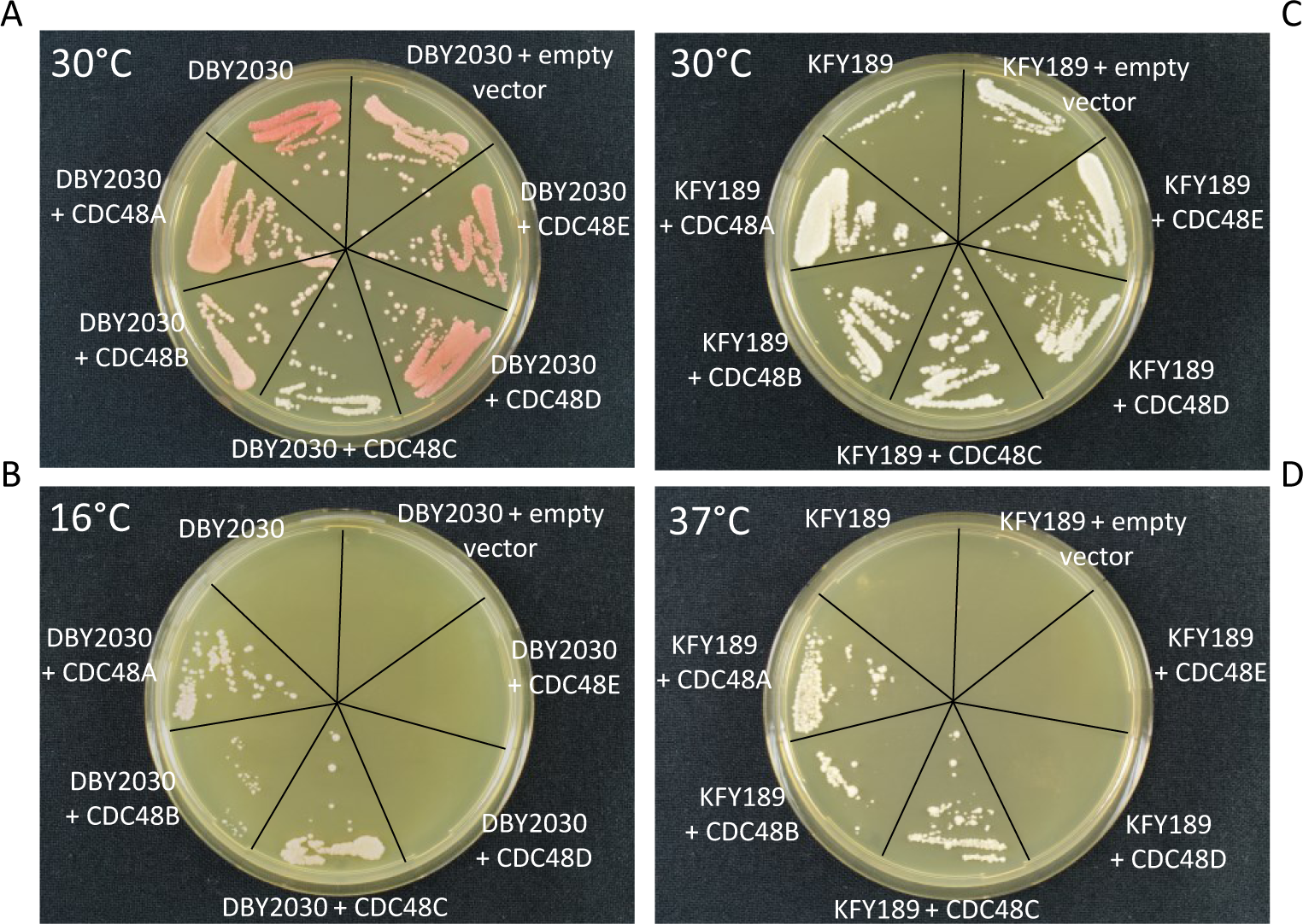
Functional complementation of yeast *cdc48* mutants. The yeast *cdc48* mutant strains which are cold-sensitive (strain DBY2030) or heat-sensitive (strain KFY189) were transformed with pDRF1-GW vector containing or not the cDNA coding Arabidopsis CDC48A-E proteins. Yeast strains were then streaked out on YPD agar plates and grown at 30°C (A, C) or 16°C for DBY2030 (B) or 37°C for KFY189 (D). Experiments were repeated three times with similar results.

Focusing on *CDC48A*, *B* and *C*, we next found that only *CDC48A* and *CDC48B* showed an increased expression in response to *B. cinerea*, at 48 and 72 hpi (Figure 5A). We then evaluated the susceptibility of different *cdc48* mutants to *B. cinerea*. We used *cdc48a-4/muse8* which is a partial loss-of-function EMS mutant previously described (36), the knock-out *cdc48b3* (GABI_104F08) and knock-down *cdc48b4* (GABI_485G04) mutants that show no or reduced *CDC48B* expression, respectively (Supplemental Figure S4A), and the two knock-out *cdc48c2* (SALK_102955C) and *cdc48c3* (SALK_123409) that do not express *CDC48C* (Supplemental Figure S4B). When compared to challenged WT plants, *cdc48a-4/muse8* and *cdc48b-4* mutants were slightly, but significantly more resistant to *B. cinerea*, as judged by the lesion size (Figure 5B, Supplemental Figure S5). *cdc48b-3* and *cdc48c-2* mutants were even more resistant, and *cdc48c-3* exhibited the strongest resistance with a 37% reduction in lesion diameter. We also produced Arabidopsis lines overexpressing either a WT AtCDC48B protein or a mutated one (AtCDC48B^E308QE581Q^ further abbreviated AtCDC48B-QQ) under the control of a CaMV35S promoter, the latter being a negative dominant allele (40). Two overexpressing lines were selected for each construct (Supplemental Figure S4C). Upon challenged with *B. cinerea*, plants overexpressing AtCDC48B-QQ were more resistant to the infection whereas plants overexpressing AtCDC48B-WT were as susceptible as WT plants (Figure 5B, Supplemental Figure S5). These data are consistent with results obtained using T-DNA insertion lines (Figure 5B), and further indicate that CDC48 activity, likely though ERAD pathway, is required for full Arabidopsis susceptibility to *B. cinerea*.

**Figure 5:**
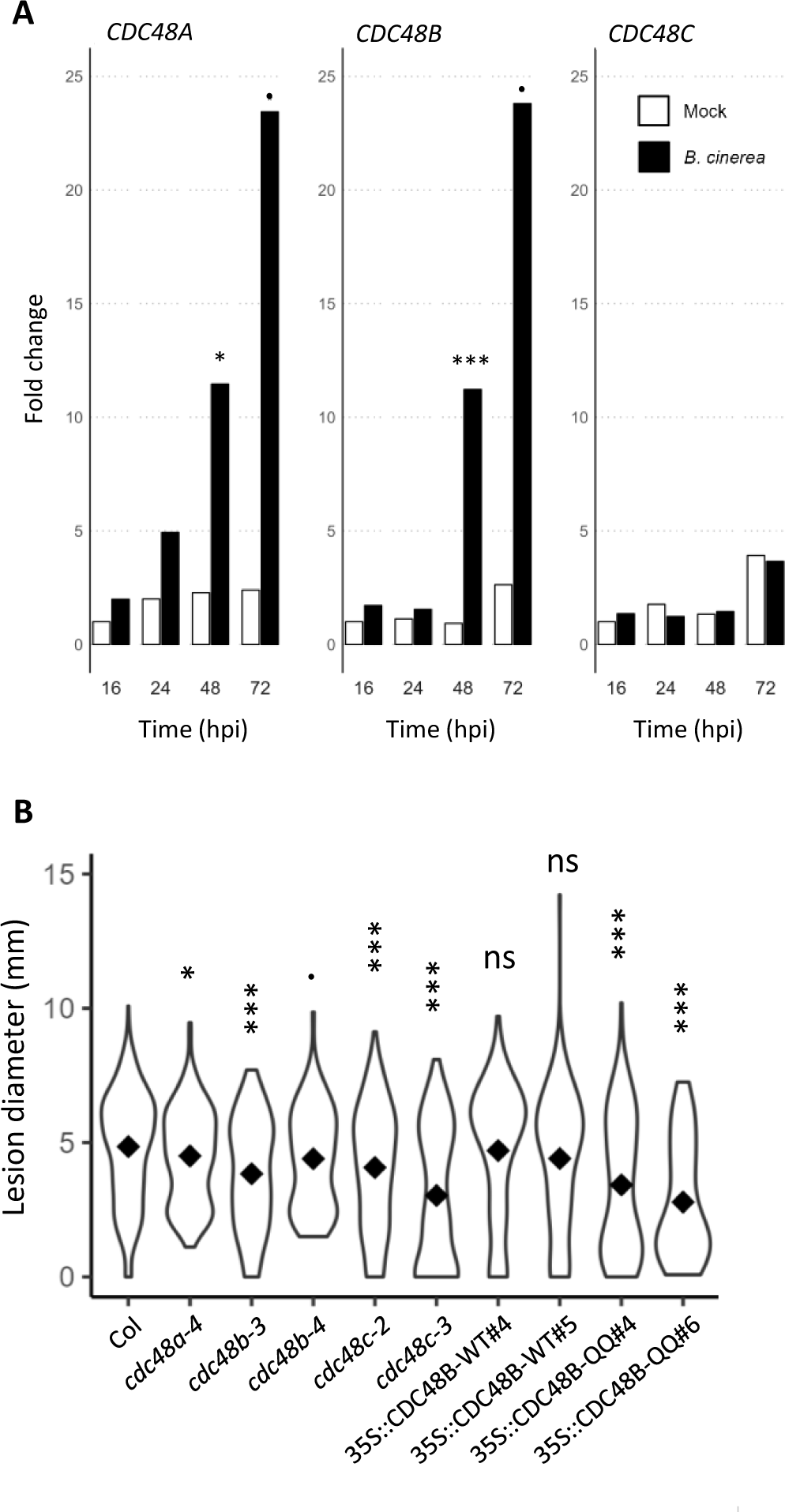
CDC48 proteins are involved in disease susceptibility caused by *B. cinerea.* (A) Expression of *CDC48A*, *CDC48B and CDC48C genes* in response to *B. cinerea* infection. Plants were sprayed with ¼ PDB solution (Mock, white bars) or with ¼ PDB solution containing *B. cinerea* spores (black bars). For each experiment, four leaves of three plants were collected at 16, 24, 48 and 72 hours post-inoculation and used for total RNAs extraction. Transcript levels were quantified by RT-qPCR and normalized to those of two reference genes, *AT4G26410* and *AT3G01150* (54). The represented fold change is the mean of 6 independent experiments. Significant differences between non-infected and infected plants were assessed by one-way ANOVA followed by a Tukey HSD post-hoc test (*** p<0.005; * p<0.05; p<0.1). (B) Disease phenotypes of *cdc48* mutants, *CDC48B-WT* overexpressors and *CDC48B-QQ* dominant negative overexpressors. Lesion diameters observed in wild-type (Col-0) and transgenics were measured 3 days after *B. cinerea* infection. Experiments were repeated seven to nine times. Seven plants per genotype were infected in each experiment. The stars indicate significant differences compared to the wild type and were assessed using Kruskal-Wallis’ method followed by Dunnett’s post-hoc test (*** p<0.005; * p<0.05; p<0.1).

Inactivation of IRE1/bZIP60 branch impacts neither the expression of *B. cinerea*-induced defence genes, nor the expression of ER-stress-induced cell death genes.

Following *B. cinerea* perception, plant cells activate a complex array of defence mechanisms that relies, at least in part, on a transcriptional reprogramming. Among genuine defence gene markers induced upon B. cinerea infection, one may cite *PR1*, *PDF1.2a*, *ORA59*, *FLG22-INDUCED RECEPTOR-LIKE KINASE 1* (*FRK1)*, *PAD3* and *GLUTHATIONE S-TRANSFERASE 6* (*GSTF6)* (Supplemental Figure S2; (24,41,42)). We reasoned that *bzip60-3 and ire1a-2 ire1b-4* mutants could have a reduced defence response if bZIP60 transcription factor positively regulates defence gene expression. To test this hypothesis, changes in the aforementioned gene expression was measured by qRT-PCR in the single and double mutants and compared to WT. In a WT genetic background, infection triggered an increase in expression for all genes at 24 and 48 hpi, except for *PR1* whose enhanced transcript level was only detected at 48 hpi (Supplemental Figure S6). This expression pattern is consistent with previous work (41). However, neither *ire1a-2 ire1b-4*, nor *bzip60-3* mutations impacted the expression profiles of the tested defence marker genes in response to *B. cinerea* (Supplemental Figure S6).

Evidence have been accumulating that necrotrophic pathogens, like *B. cinerea*, actively induce host PCD for feeding on cell debris and subsequent colonization of dead tissues (27). With this regard, we hypothesized that ERSID could be over-induced in the *bzip60-3 and ire1a-2 ire1b-4* mutants, potentially explaining their enhanced susceptibility to the fungus (Figure 1A). ERSID was found to be positively controlled by the NAC089 transcription factor (10) and the protease family of cathepsins (11) under chemically-induced ER stress. Whereas *NAC089* and *CATHEPSIN B1, B2, B3* gene expression was previously shown to be significantly increased during ERSID triggered by tunicamycin (10,11), none of those genes were differentially expressed between mock and infection (24 and 48 hpi) conditions in a WT genetic background (Supplemental Figure S7). In addition, no or minor expression changes for those genes could be recorded in *bzip60-3 and* double *ire1a-2 ire1b-4* mutants compared to WT, whether plants were inoculated or not (Supplemental Figure S7).

The expression of NAC053, a negative regulator of Arabidopsis defence against *B. cinerea*, is dependent on IRE1 proteins.

To explain the susceptibility phenotype of *ire1a-2 ire1b-4* and *bzip60* mutants and the resistant phenotype of *cdc48* mutants, we tested the hypothesis that the IRE1/bZIP60 branch of the UPR negatively control *CDC48* gene expression and its regulators in response to *B. cinerea*. None of the *CDC48A, B, C* genes were mis-regulated at 24hpi and 48hpi in *bzip60-3* and *ire1a-2-ire1b-4* mutants (Supplemental Figure S8). The expression of the two NAC TF (NAC053/NTL4 and NAC078/NTL11) that have been shown to regulate proteasome stress regulon-encoding genes, including *CDC48A* (Gladman *et al.*, 2016), was next checked in the same genetic backgrounds (Figure 6). The expression level of *NAC078* was not altered by the infection in any of the genotypes for the two time points tested (Figure 6A-B). By contrast, in response to *B. cinerea*, *NAC053* expression was induced earlier in the *bzip60-3* mutant (24hpi, Figure 6A) when compared to WT (48hpi, Figure 6B), and was totally abolished in the double *ire1a-2-ire1b-4* mutant at both time points (Figure 6A-B). Infected *nac053-1* and *nac078-1* mutants showed smaller lesions at 3 dpi with respect to WT plants (Figure 6C, Supplemental Figure S9), indicating that both mutants are more resistant to *B. cinerea*. These results indicate that *NAC053* expression is dependent on IRE1 proteins, and that both NAC053 and NAC078 act as negative regulator of Arabidopsis defence against *B. cinerea*.

**Figure 6:**
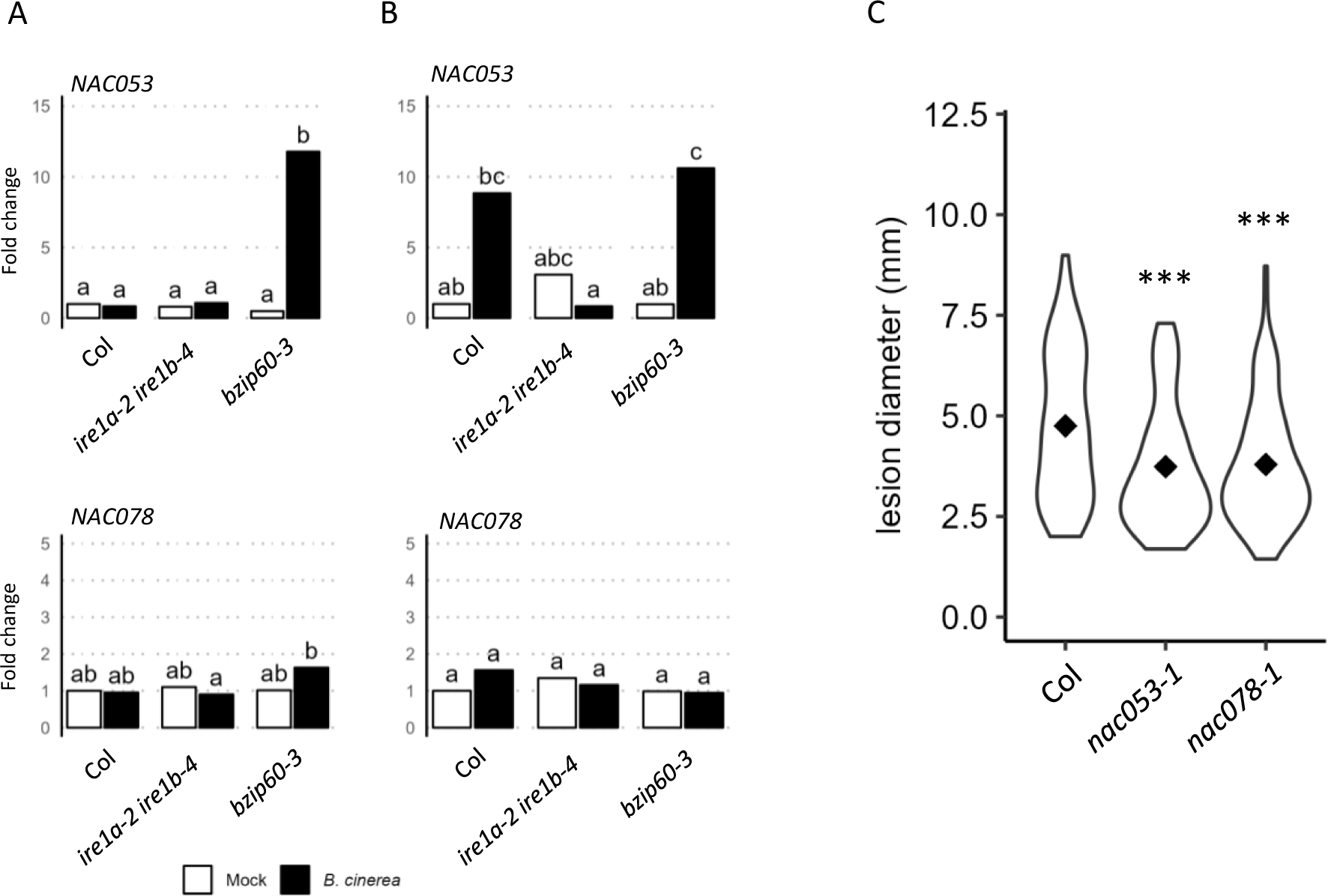
Defect in NAC053 and NAC078 transcription factors reduces *B. cinerea-* induced symptoms. (A-B) Expression of *NAC053* and *NAC078* genes in response to *B. cinerea* infection in WT, *ire1a/ire1b* and *bzip60* mutants. Plants were sprayed with ¼ PDB solution (Mock, white bars) or with ¼ PDB solution containing *B. cinerea* spores (black bars). For each experiment, four leaves of three plants were collected at 24 h (A) or 48 h (B) post infection and used for total RNA extraction. Transcript levels were quantified by RT-qPCR and normalized to those of two reference genes, *AT4G26410* and *AT3G01150* (54). The represented fold changes are the mean of six independent experiments. Different letters represent groups which were significantly different from one another as determined by a one-way ANOVA followed by a multiple comparison with a Fisher’s Least Significant Difference (LSD) test. (C) Disease phenotype of *nac053* and *nac078* mutants infected with *B. cinerea*. Lesion diameters observed in wild-type (Col) and mutants were measured 3 days after *B. cinerea* infection. Experiments were repeated four times. Seven plants per genotype were infected in each experiment. The stars indicate significant differences compared to the wild type. Significant differences between a mutant genotype and WT plants were assessed using Kruskal-Wallis’ method followed by Dunnett’s post-hoc test (*** p < 0.005; ns: non-significant difference).

## Discussion

The UPR pathway plays a crucial role in plant-pathogen interactions, irrespective of the pathogen nature, *i.e.* bacteria, fungi and viruses (13,44). However, our understanding of its specific role during plant infection by necrotrophic fungi remains elusive. Here we present a study that identifies UPR pathway as a critical element in *Arabidopsis thaliana* response to two necrotrophic fungi, *Botrytis cinerea* and *Alternaria brassicicola*. The RT-qPCR results demonstrated that *B. cinerea* infection activated UPR pathways in WT plants. During infection, we observed an accumulation of mRNAs coding the three BIP isoforms, along with its co-chaperones SDF2.2, ERDJ3A and ERDJ3B. Additionally, we detected an increased level of spliced bZIP60 mRNA variants coding the active TF. The importance of UPR has been confirmed by challenging plants mutated in the genes coding the canonical UPR regulators IRE1A/IRE1B, bZIP60, as well as those of RIP-UPR bZIP17 and bZIP28, with a spore solutions of *B. cinerea*. Mutation in the genes coding *IRE1A* and *IRE1B* or *bZIP60* results in a greater plant susceptibility to *B. cinerea.* Similar observations were made when these mutants were challenged with *Alternaria brassicicola* indicating that IRE1-bZIP60 pathway plays important role in limiting the growth of both pathogens. However, it is worth noting that mutation in *bZIP28* did not have a significant impact on the plant susceptibility to these fungi. Overall, our findings support the hypothesis that the IRE1-bZIP60 branch is an important element in the establishment of plant immunity to necrotrophic pathogens. This is in accordance with previous work on *Nicotiana benthamiana*, where silencing of *NbbZIP60*, *NbIRE1a*, and *NbIRE1b* expression through a VIGS approach led to increased plant susceptibility to the necrotrophic fungus *Alternaria alternata* (14). Furthermore, infection of the double mutant *bzip28-2-bzip60-1* with *Drechslera gigantea*, a necrotrophic fungus causing eyespot disease in crop plants, leads to increase disease symptoms compared to WT plants in Arabidopsis (15). However, it is noteworthy to mention that neither the knock down *bzip60.1* nor the double mutant *bzip28-2-bzip60-1* showed increased susceptibility when infected with *B. cinerea* (Supplemental Figure S10). The significance of the IRE1-bZIP60 branch of the UPR in controlling plant immunity is further demonstrated by the work of Tateda *et al.* (45), who showed that silencing of *IRE1a/1b* or *bZIP60* resulted in disease symptoms in *Nicotiana tabacum* caused by the non-host bacterial pathogen *Pseudomonas cichorri*. Additionally, concerning the virulent bacteria *Pseudomonas syringae pv maculicola* ES4326, bacterial growth was substantially more pronounced in *ire1a ire1b* or *bzip60* infected mutants compared to the infected WT plants (46). It is must be pointed out that our study revealed distinct responses to *B. cinerea* and *A. Brassicicola* depending on the genotype tested. Notably, the *bzip17* mutant demonstrated higher resistance to *B. cinerea* compared to WT plants, while it exhibited a similar level of infection as WT plants when exposed to *A. brassicicola*. Studies demonstrating a role for bZIP17 during biotic stresses are scarce. One may cite the study of Li (47) where the silencing of both *NbbZIP17* and *NbbZIP28* delayed Rice Stripe Virus infection and decreased the accumulation of viral RNA and proteins in *N. benthamiana*, indicating that RIP-UPR promotes virus development. However, in Arabidopsis infected with the *Plantago asiatica* mosaic virus, bZIP60 and bZIP17 acts synergistically to restrict viral infection (48). Understanding the role of bZIP17 during *B. cinerea* infection will need further investigations.

Activation of UPR leads to an increase of the ER folding capacity, preventing an excessive build-up of misfolded or aberrant proteins that may occur during stressful situations like pathogen attacks. Unfolded and misfolded proteins can also be degraded through ERAD. During this process, misfolded proteins are tagged with polyubiquitin chains and extracted from the ER membrane with the assistance of the CDC48 AAA+ATPase complex (9). The number of gene coding CDC48 proteins in Arabidopsis genome is unclear (35). Through yeast *cdc48* mutant complementation experiments, our findings unequivocally establish that only three proteins - AtCDC48A (At3g09840), AtCDC48B (At3g53230), and AtCDC48C (At5g03340) - are functional orthologs of ScCDC48 (as demonstrated in our study and by Feiler *et al.* for CDC48A (38)). Thus, the two closely related AAA+-ATPase proteins (At2g03670 and At3g01610), previously referred to as AtCDC48D and E (35,36), are not ScCDC48 functional orthologues and should not be categorized as such.

An increase in the accumulation of *AtCDC48A* and *AtCDC48B* mRNA have been detected during *B. cinerea* infection. Interestingly, plants mutated in either *AtCDC48A*, *AtCD48B* or *AtCDC48C* genes are more resistance to *B. cinerea* infection. Furthermore, the overexpression of an inactive form of AtCDC48B (*eg* AtCDC48B^QQ^) also results in an increased resistance phenotype toward *B. cinerea*. The resistant phenotype of *atcdc48a-4* mutant to *B. cinerea* could be tentatively explained by a constitutive immune response characterized by the stronger accumulation of *PR2* mRNA in this mutant (36). Ao et *al.* (49) have proposed a model to explain this autoimmune phenotype related to the role of AtCDC48A during plant pathogen interaction. AtCDC48A is recognized by the SFC^SNIPER7^ complex and targeted for proteasomal degradation. It results in the accumulation of the NLR SNC1 (and likely other NLR) which may serve to enhance defence to the virulent oomycete pathogen *H. arabidopsidis* NOCO2. The existence of such a mechanism in the context of *B. cinerea* infection remains to be demonstrated. Regarding AtCDC48A paralogs, AtCDC48B and AtCDC48C, it is unlikely that they have redundant functions with AtCDC48A. This is supported by the observations that mutation in each of the *AtCDC48* gene results in resistant phenotypes against *B. cinerea.* Nonetheless, mutations in *AtCDC48B* or *AtCDC48C* do not affect the resistance of Arabidopsis to *H. Arabidopsis* NOCO2 (36). However, the importance of the IRE1-bZIP60 pathway in basal resistance to *B. cinerea* infection is unlikely linked to CDC48 because *AtCDC48* mRNAs do not accumulate differentially in *atire1a/ire1b* or *atbzip60* genetic backgrounds. Understanding the involvement of each of the *AtCDC48* genes in plant immunity will necessitate additional in-depth investigations.

To identify potential mechanisms that play a role in the defence response against *B. cinerea* via the IRE1-bZIP60 pathway, we have undertaken a RT-qPCR approach. None of the tested genes had their expression altered in the *ire1a-2 ire1b-4* and *bzip60-3.* It concerns the SA (*PR1*) and JA (*PDF1.2, ORA59*) responsive genes, indicating that the IRE1-bZIP60 pathway does not influence the expression of SA and JA-dependant gene expression in response to *B. cinerea* infection. However, we did not investigate the hypothesis whether SA or JA signalling pathways were necessary to regulate the activation of UPR pathways as observed in other studies. In tomato, JA treatment induced the expression of *BIP1* and tunicamycin-induced *BIP1* expression was strongly reduced in the JA signalling mutant *jai1* indicating that JA pathway plays a role in ER stress signalling (50). Similar observations were made in *N. attenuata* in which MeJA treatment induced the accumulation of mRNA coding chaperone proteins such as BIP, PDI, CNX and CRT as well as the spliced form of *bZIP60* mRNA (14). Using JA-deficient and JA-insensitive plants, these authors have also shown that JA controls chaperone gene expression in response to *A. alternata*.

In addition, our data indicate that *PAD3* gene, which encode a cytochrome P450 enzyme that catalyses the last step of camalexin biosynthesis is not regulated by UPR. To the contrary, the *F6’H1* gene was found less induced in *N. attenuata* plants impaired in the IRE1-bZIP60 pathway during *A. alternata* infection (14). It encodes the key enzyme which catalyzes the synthesis of scopoletin and scopolin, two phytoalexin previously shown to be crucial for *N. attenuata* resistance to *A. alternata*. These results indicate that the role of UPR in regulating plant resistance to one particular class of pathogen is different from one plant-pathogen interaction model to another.

Necrotrophic pathogens induce plant cell death to their own benefits either by secreting plant cell death inducers or by manipulating host regulated cell death (27,28). Genes known to orchestrate ERSID, such as *NAC089* and *CATHEPSIN B*, are not induced after *B. cinerea* infection whereas these genes are up-regulated following plant treatment with the ER stress-inducing agent tunicamycin (10,11). We can formulate two hypotheses. Firstly, increased susceptibility of *ire1a-2 atire1b-4* and *atbzip60-3* to *B. cinerea* is unlikely the consequence of an increase ERSID phenomenon because *NAC089* or *Cathepsin B* genes are not upregulated in these mutant genotypes. It suggests that a NAC089- and CATHEPSIN B-dependant ERSID is unlikely to be involved during Arabidopsis*-B. cinerea* interaction. Secondly, we can hypothesized that ER stress is not involved in *B. cinerea* - induced cell death. In any case, further experiments are needed to clarify the role of ER stress in *B. cinerea*-induced cell death.

Our data indicate that, in *bzip60* mutant infected with *B. cinerea*, the expression of NAC053/NTL4 is upregulated. Infected-*bzip60* mutant accumulates much more *NAC053* mRNA than WT plant and this accumulation occurs earlier in *bzip60* (at 24 hpi) than in WT plant (at 48 hpi). However, the expression of the NAC053 closest relative NAC078/NTL11 is not modulated by *B. cinerea* in any of the genotypes tested. Together, both transcription factors are required to control the expression of the proteasome stress regulon including *CDC48A* (43). Nevertheless, the expression of genes coding CDC48 A, B or C proteins are not modulated in Botrytis-infected *bzip60* mutant. Very surprisingly, results are different when using *ire1a-ire1b* double mutant. *NAC053* mRNA does not accumulate neither at 24 hpi nor at 48 hpi in this mutant infected with *B. cinerea* in comparison to WT plants in which *NAC053* accumulate at 48 hpi. It indicates that *NAC053* expression depends on IRE1 proteins during *B. cinerea* infection. NAC053 is anchored to the plasma membrane (51) and activated following stress perception in a ROS-dependant mechanism (52). Once activated, NAC053 control ROS production through the activation of *RBOH* genes (53). In our model, it is reasonable to suggest that a disturbance in ROS homeostasis is likely present in the *bzip60* mutant, whereas such a disturbance might not exist in the *ire1a/ire1b* mutant. Further research is necessary to substantiate this hypothesis.

Summarizing, the infection of wild type Col-0 plant by *Botrytis cinerea* induces ER stress and activates the UPR pathways (Figure 7). The activation of the IRE1-bZIP60 branch and ERQC is crucial for the plant’s response to the pathogen, as mutations in genes encoding key players in these pathways result in increased plant susceptibility. Further work is however necessary to understand how the IRE1/bZIP60 branch contributes to defence against this necrotrophic fungus, particularly its potential involvement in the secretion of defence proteins. On the other hand, our results indicate that mutations in the bZIP17 branch of the UPR, as well as mutations in genes encoding the ERAD-involved proteins CDC48, lead to greater resistance of the plant. This suggests either that these proteins act as negative regulators of immunity against this pathogen, or that these pathways might be manipulated by the pathogen during infection. Similarly, further work is needed for a detailed understanding of these mechanisms.

**Figure 7:**
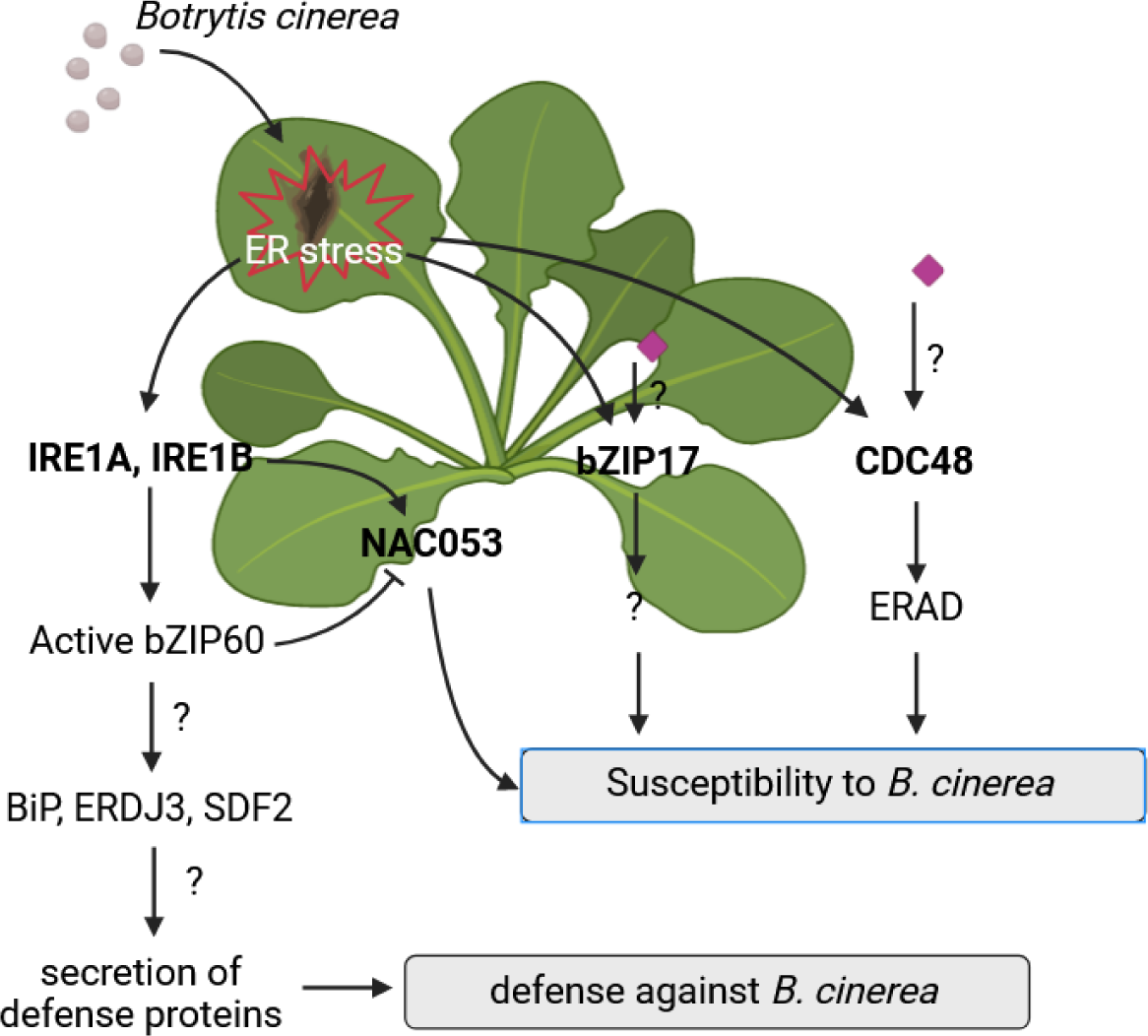
Schematic representation of the role of UPR, ERQC and ERAD during *Botrytis cinerea* infection of *Arabidopsis thaliana* plants. *Botrytis cinerea* infection induces an ER stress which activates the canonical IRE1-bZIP60 branch of the UPR and the expression of the genes coding the ERQC machinery (BIP, ERDJ3, SDF2). We hypothesized the activation of these signalling pathways might control the secretion of defence proteins which are necessary to establish defence against *B. cinerea*. Our data also indicate that active bZIP60 suppresses *NAC053* expression, a negative regulator of defence against *B. cinerea*. However, *NAC053* expression depends on IRE1. Regarding the bZIP17 arm of the UPR or the ERAD proteins CDC48, we hypothesized that they act as negative regulators of immunity against B. cinerea or might be the target of *B. cinerea* effectors (purple diamonds) thus facilitating infection process.

## Experimental procedures

### Plant materials

*A. thaliana* seeds were grown on jiffy-7 pellets in a growth chamber under controlled conditions (10h light/14h dark, 22°C/18°C, 60–70% humidity, light intensity of 100 and 120 *μ*E.m^−2^.sec^−1^). All the mutants used are listed in Supplemental table S1. To screen T-DNA homozygous plant mutants by PCR, genomic DNA was isolated using Phire Plant Direct PCR kit (Thermoscientific) and PCR was performed using primers pairs as indicated in Supplemental table S2.

### Pathogen infection

*Botrytis cinerea* strain BMM (29) was grown 10 days on sterile V8 agar plate (50% Campbells original V8 juice (v/v), 0.5 % KH2PO4 (m/v), 1.5 % agar (m/v), pH 6) at 20°C in the dark. Spores were harvested by scraping the plate with sterile milliQ water and filtered through sterile gaze. Infection tests are carried out by placing 6 µL of a solution of botrytis spores (5.10^4^ spores.mL^−1^ diluted in quarter-strength Difco potato dextrose broth PDB) on 4 leaves of four weeks old plants. Lesion diameters were measured after 3 days. For qPCR analysis, plants were either sprayed with a spore solution (2.5.10^5^ spores.mL^−1^) or with quarter-strength PDB as described previously (41). For both experiments, the inoculated plants are placed in a tray closed with a transparent lid to maintain a high-humidity environment.

*Alternaria brassicicola* strain MIAE01824 was provided by Dr. Christian Steinberg (INRAe, Dijon, France) and grown 15-20 days on PDA medium (19 g.L^-1^) supplemented with sucrose (20 g.L^-1^) and CaCO (30 g.L^-1^) at 20°C in the dark. Spores were harvested by scraping the plate with sterile infection medium (GamborB5 medium (Duchefa), sucrose 10 mM, KH_2_PO_4_ 10 mM) and filtered through sterile gaze. Four leaves of four weeks old plants were drop inoculated (6 µL) with a solution of 1.10^6^ spores.mL^−1^ in infection medium. Inoculated plants are placed in a tray closed with a transparent lid to maintain a high-humidity environment and lesion diameters were scored after five days.

### RNA extraction and RT-qPCR analysis

Inoculated and mock-treated leaves were harvested at different time points, flash frozen and finely grinded in liquid nitrogen. Total RNA extraction and DNAse treatment were performed using SV Total RNA Isolation System (Promega) as described by the manufacturer. RNA integrity was analysed on a 1% agarose gel and concentration was measured using a nanodrop system (Thermoscientific). RNA (500 ng) were retrotranscribed into cDNA (High-Capacity cDNA Reverse Transcription Kit, Thermoscientific) using random hexamer and 17-mer oligodT primers. mRNA accumulation was assessed by real-time qPCR (GoTaq® qPCR Master Mix, Promega) and expression values were normalized to the expression of plant genes *At4g26410* and *At3g01150* as previously described as a stable reference genes (54). Levels of transcripts were calculated using efficiency-weighted ΔΔCq^ω^ method (55). All primers are listed in Supplemental table S3.

### Statistical methods

For infection tests, significant differences from WT plant were determined by a Kruskal–Wallis one-way anova on ranks followed by a comparison with the Dunnett’s method. For qPCR analysis, ΔCq^ω^ value from one infection time-point is compared to the data of the control condition at the same time point using by a one-way ANOVA followed by a Tukey HSD Test (55). Then, data are represented on the figures as fold induction to the control condition (ΔΔCq^ω^).

### AtCDC48B cloning procedures and transgenic lines generation

Arabidopsis lines overexpressing either AtCDC48B or AtCDC48B mutated in its two ATPase sites (AtCDC48B^E308QE581Q^) under the control of a CaMV35S promoter were produced as follows. PDONR/Zeo containing either AtCDC48B^WT^ or AtCDC48B^E308QE581Q^ cDNA were kindly provided by Drs. Annette Niehl and Manfred Heinlein (IBMP, Strasbourg, France). pDNOR/Zeo were BP-recombined (Thermoscientific) in pB2GW7 vector (56) and resulting vectors were introduced in *Agrobacterium tumefaciens* strain GV3101. WT *A. thaliana* Columbia plants (N60,000, Eurasian Arabidopsis Stock Center) were transformed using the floral dipping method (57). T1 and T2 transgenic plants were grown on jiffy-7 pellets and selected using BASTA herbicide after 10 days of growth. To identify homozygous plants overexpressing *AtCDC48B*^WT^ or *AtCDC48B*^E308QE581Q^, seeds from T2 plants were surface-sterilized and sown on MS Agar plates (4.4 g.L^-1^ of Duchefa MS powder M0222.0050, 1% sucrose, 10 mM MES, 1% agar pH 5.7) containing 50 µg/mL of glufosinate ammonium (Sigma-Aldrich). Expression of mRNA coding AtCDC48B^WT^ or AtCDC48B^E308QE581Q^ in transgenics plants were analysed by RT-qPCR using primers described in supplemental table S3.

### CDC48 expression in *cdc48* yeast mutant

CDC48 cDNA were cloned from an Arabidopsis thaliana cDNA pools made from total RNA extracted from Arabidopsis Col-0 plants. AtCDC48A (At3g09840; forward primer ATGTCTACCCCAGCTGAATCTTC; reverse primer CTAATTGTAGAGATCATCATCGTCCC), AtCDC48C (At5g03340; forward primer ATGTCAAACGAACCGGAATC; reverse primer CTAACTGTAGAGATCGTCGTCATC), AtCDC48D (At2g03670; forward primer ATGTTGGAAACCGAAAGC; reverse primer TCATGTAGCAGAAGCTACTAGTAA), AtCDC48E (At3g01610; forward primer ATGGGGAGGAGAGGTCGC; reverse primer TTACTCGAGGGTAAAAGATGGCC) were amplified by PCR using Phusion DNA polymerase (Thermoscientific) and cloned into PCR8 vector (Thermoscientific). PDONR/Zeo containing CDC48B (At3g53230) was kindly provided by Drs. Annette Niehl and Manfred Heinlein (Institut de biologie moléculaire des plantes, Strasbourg, France; (40)). PCR8 and pDNOR/Zeo vectors were then recombined in pDRF1-GW (58) and resulting vectors or empty vector were introduced in *cdc48* deficient *Saccharomyces cerevisiae.* Two *cdc48 S. cerevisiae* mutants (39) were kindly provided by K.-U. Fröhlich (University of Graz, Austria): the cold-sensitive strain DBY2030 (MATa ade2-101 lys2-801 ura3-52 cdc48-1;) or the temperature-sensitive strain KFY189 (MATa lys2 leu2 ura3 cdc48-8). Transformed *S. cerevisiae* cells were plated on minimal selective medium containing 2% glucose, 0,77g/L SD-URA drop out mix (Clontech), 0.67 g/L Yeast Nitrogen Base (Sigma-Aldrich), 15 g/L agar (Sigma-Aldrich) and grown at 30°C. Transformed colonies were checked by colony PCR and then grown overnight at 30°C under shaking in YPD medium (10 g/L yeast extract, 20 g/L peptone, 2% glucose). Yeast cells were then streaked out on YDP agar plate (YPD medium containing 15g/L agar) and grown at 30°C and 16 °C for DBY2030 transformed strain or 30°C and 37°C for KFY189 transformed strain.

## Acknowledgments

We would like to thank all the people who provided us with invaluable assistance in carrying out this work and particularly Pr. Nathalie Leborgne-Castel, Dr. Mathieu Gayral, Dr. Stéphane Bourque and Pr. David Wendehenne for helpful discussions. Pr. Steven H Howell, Pr. Cyril Zipfel and Pr. Xin Li are kindly thanked for sharing Arabidopsis mutants with us, Dr Kai-Uwe Fröhlich for providing Yeast strains, and Dr. Christian Steinberg for giving us *Alternaria brassicicola* strain MIAE01824.

---

**Supplemental Figure S1.**
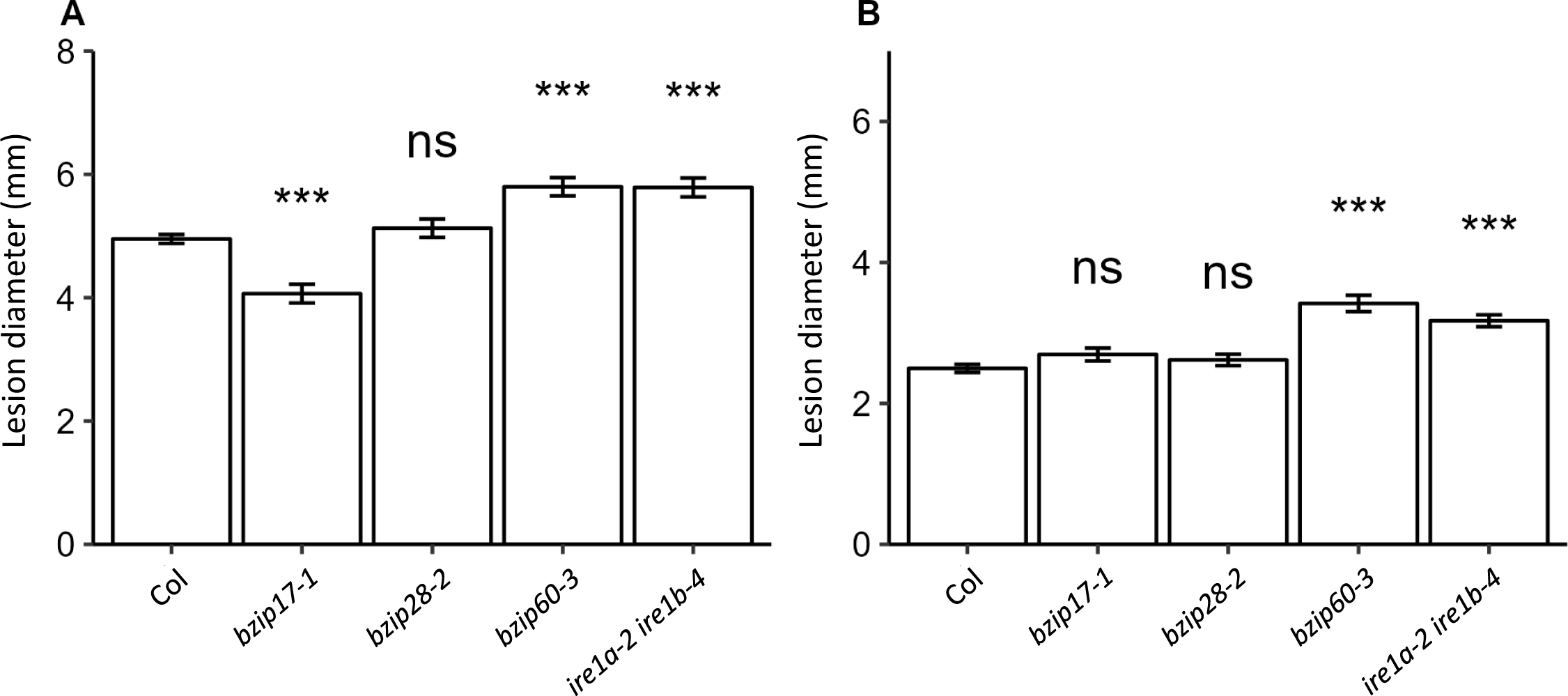
Mean lesion diameters observed on UPR-deficient mutants infected by *B. cinerea* or *A. brassicicola*. Lesion diameters were measured 3 days after *B. cinerea* infection (A) and 5 days after *A. brassicicola* infection (B). Experiments were repeated seven to eight times for *B. cinerea* infection depending on the genotype tested and four times for *A. brassisicola* infection. Seven plants per genotype were infected in each experiment. The stars indicate significant differences compared to the wild type which were identified using a Kruskal-Wallis’ method followed by Dunnett’s post-hoc test (*** p < 0.005; ns: non significant difference).

**Supplemental Figure S2:**
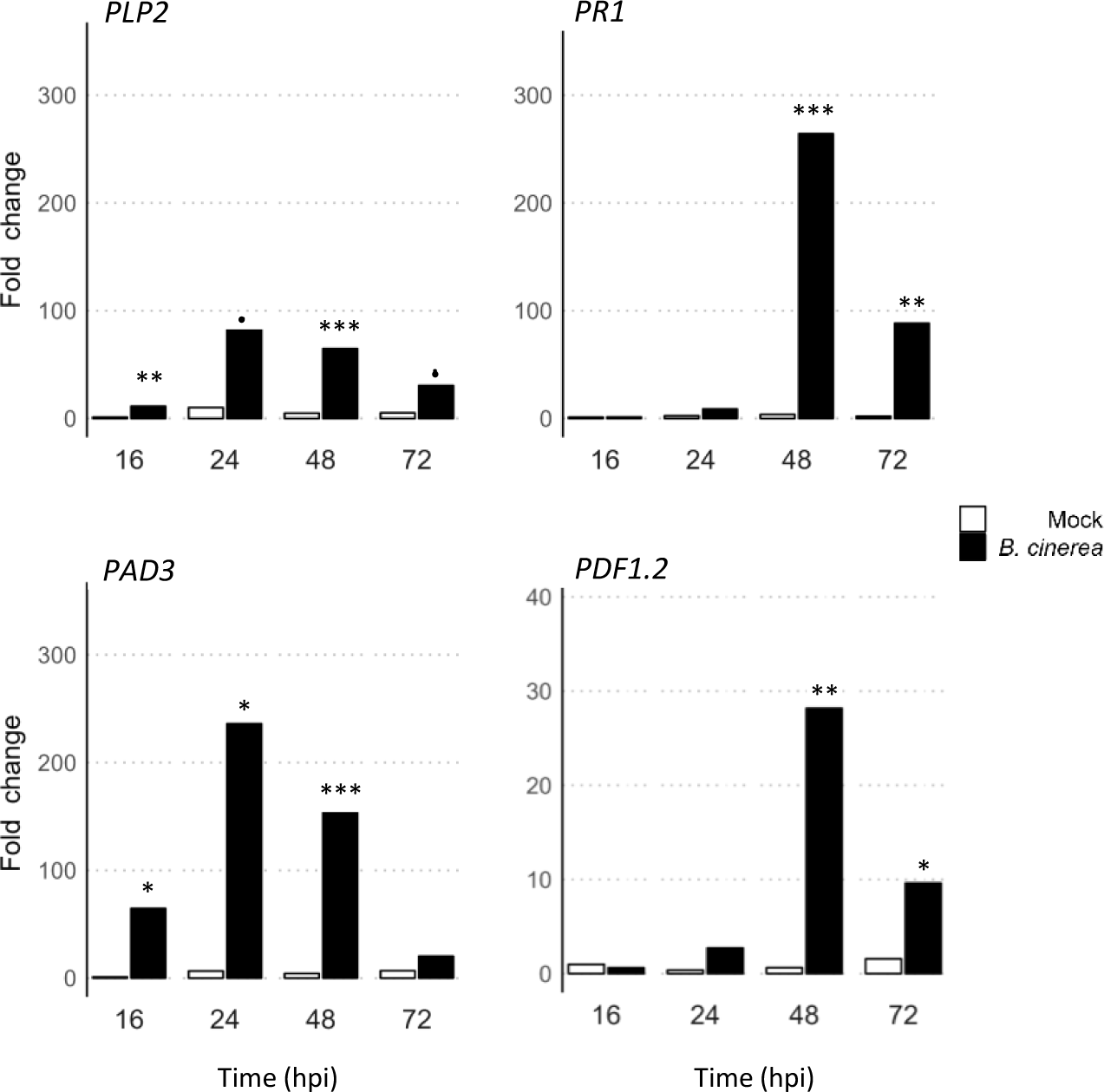
Expression kinetic of the classical Botrytis-induced defense genes *PLP2*, *PR1*, *PAD3* and *PDF1.2* in our experimental conditions. Plants were sprayed with ¼ PDB solution (Mock, white bars) or with ¼ PDB solution containing *B. cinerea* spores (black bars). For each experiments, four leaves of three plants were collected at 16, 24, 48 and 72 hours post-inoculation. Transcript levels were quantified by RT-qPCR and normalized to the plant reference genes *AT4G26410* and *AT3G01150* transcript levels (Czechowski et al., 2005). The represented fold change is the mean of five independent experiments. Significant differences between non-infected and infected plants were determined by a one-way ANOVA followed by a Tukey HSD post-hoc test (*** p<0.005; ** p<0.01; * p<0.05; • p<0.1).

**Supplemental Figure S3:**
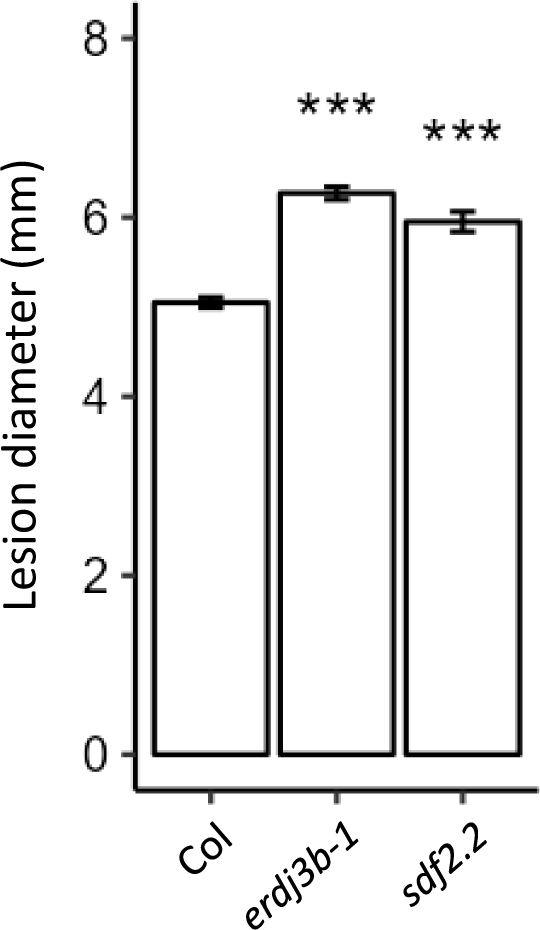
Mean lesion diameters observed on ERQC mutants infected by *B. cinerea*. Lesion diameters observed on wild-type Col plants and ERQC mutants were measured 3 days after *B. cinerea* infection. Experiments were repeated three to four times. Seven plants per genotype were infected in each experiment. The stars indicate significant differences compared to the wild type and were identified using Kruskal-Wallis’ method followed by Dunnett’s post-hoc test (*** p < 0.005).

**Supplemental Figure S4:**
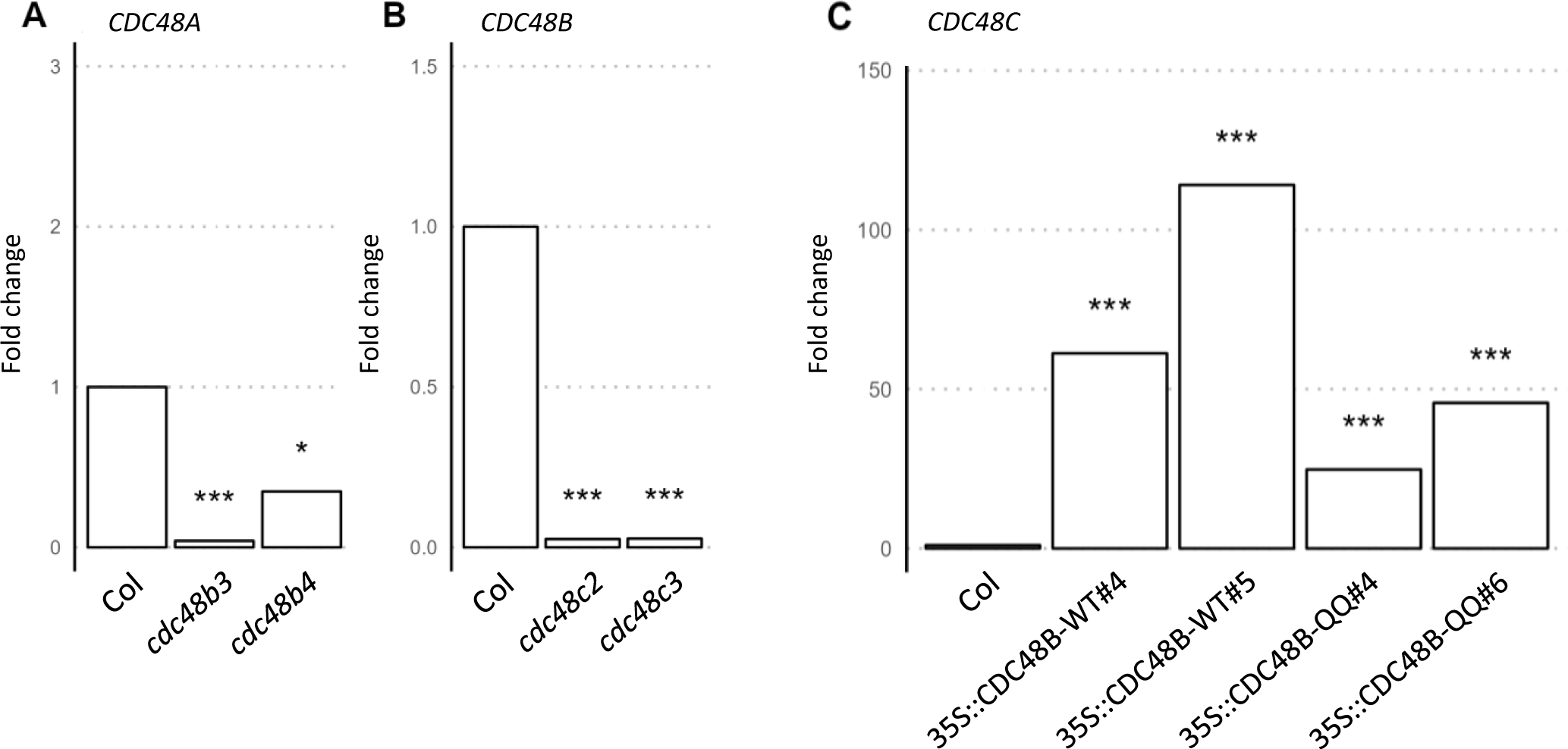
Expression of *CDC48* in *cdc48* mutants. A, B, C. Expression of CDC48 mRNAs in *cdc48b* and *cdc48c* mutants and in CDC48B overexpressing lines. Four leaves of three plants were collected and *CDC48B* or *CDC48C* transcript levels were quantified by RTqPCR in *cdc48b3* and *cdc48b4* (A) or in *cdc48c2* and *cdc48c3* (B) mutants, respectively. *CDC48B* transcript level was quantified in transgenic lines overexpressing CDC48B-WT or CDC48B^E308QE581Q^ (CDC48-QQ). All transcript level were normalized to the plant reference genes *AT4G26410* and *AT3G01150* transcript levels (Czechowski et al., 2005). The represented fold change is the mean of five independent experiment. Significant differences between non-infected and infected plants were determined by a one-way ANOVA followed by a Tukey HSD post-hoc test (*** p<0.005; * p<0.05).

**Supplemental Figure S5:**
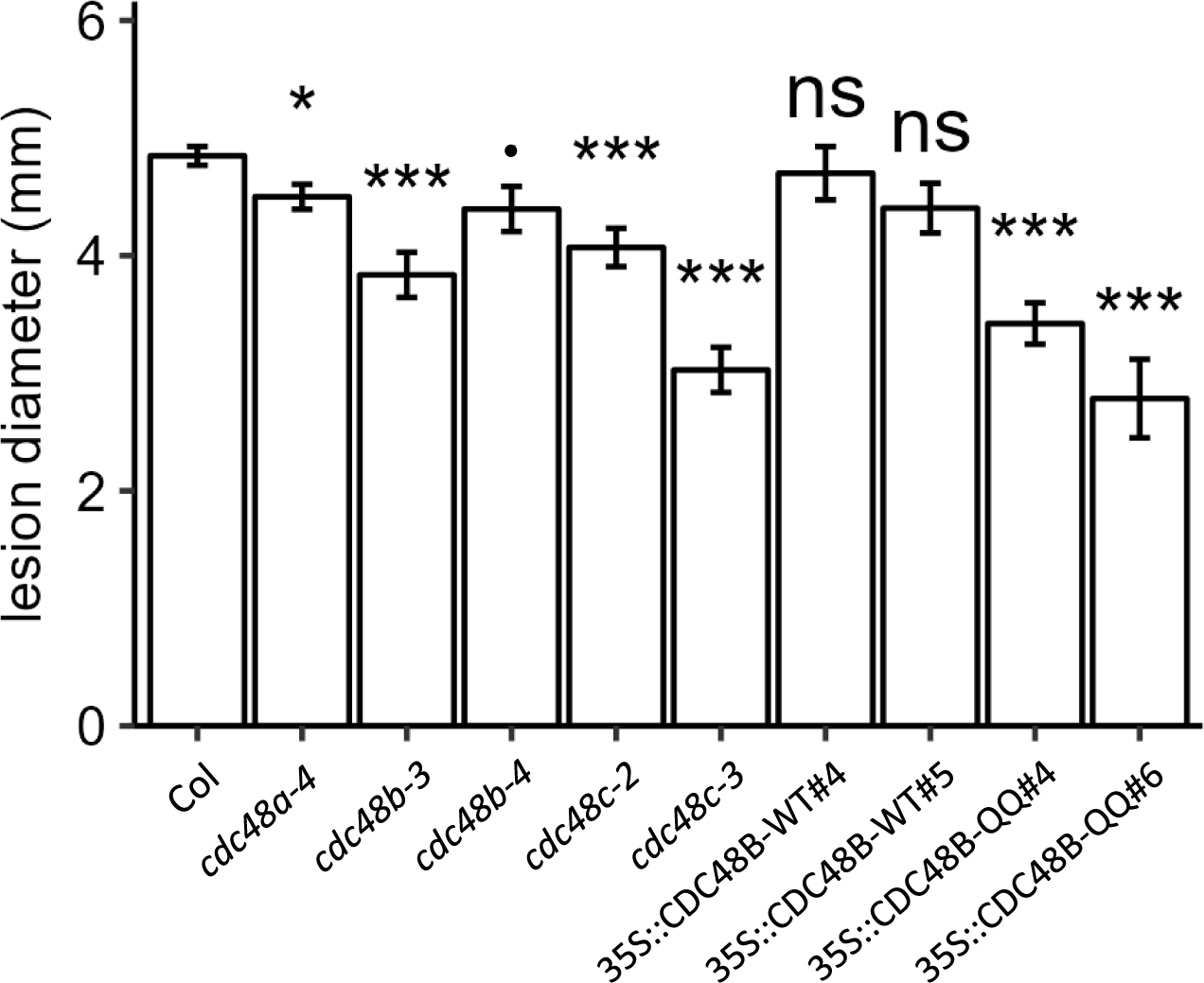
Mean lesion diameters observed on CDC48 mutants and plants expressing a WT (CDC48B-WT) or an inactive isoforms of CDC48B (CDC48B-QQ) infected by *B. cinerea*. Lesion diameters observed on wild-type Col plants and mutants were measured 3 days after *B. cinerea* infection. Experiments were repeated seven to nine times. Seven plants per genotype were infected in each experiment. The stars indicate significant differences compared to the wild type which were identified using Kruskal-Wallis’ method followed by Dunnett’s post-hoc test(*** p<0.005; * p<0.05; • p<0.1).

**Supplemental Figure S6.**
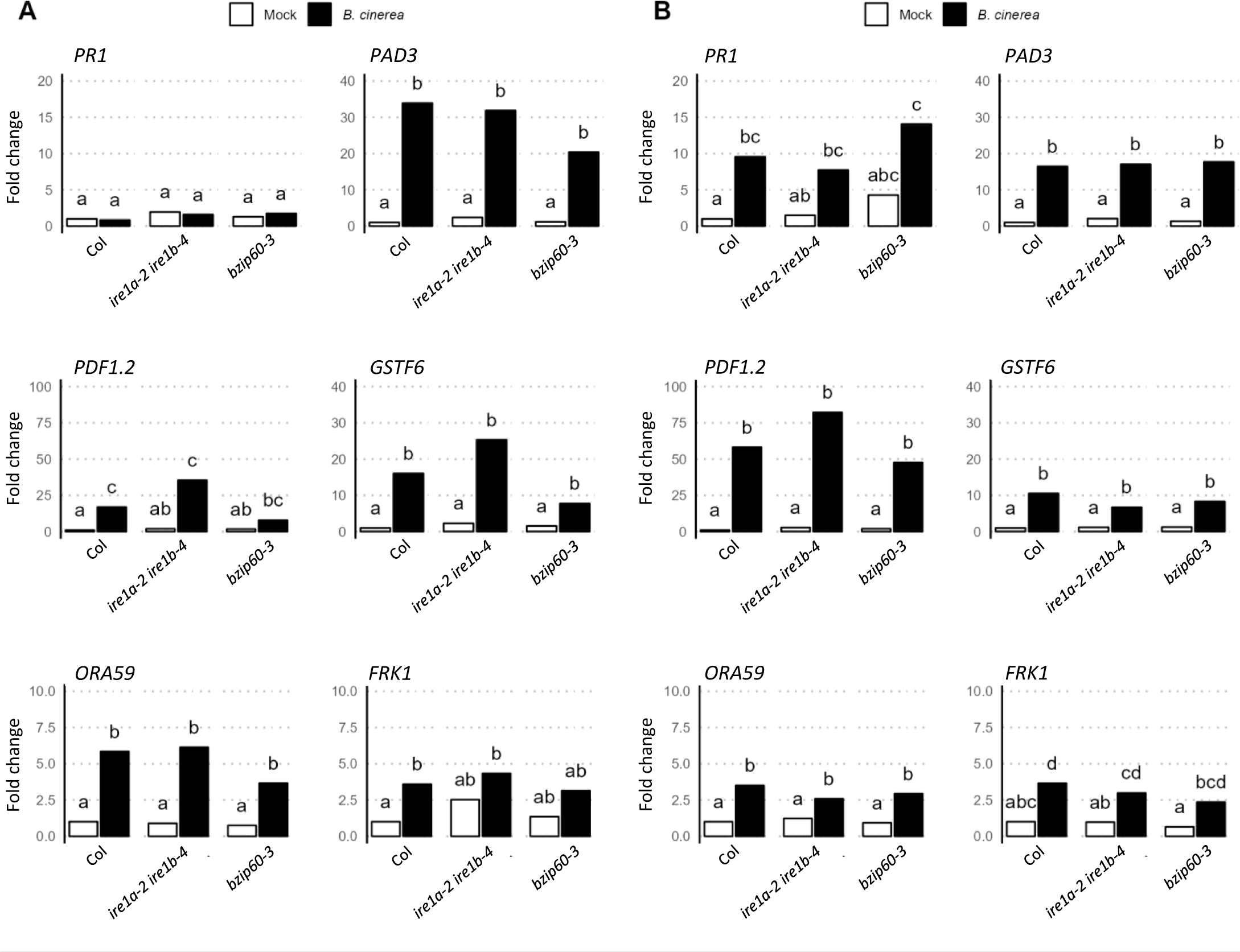
Accumulation of mRNA coding classical defense genes in response to *B. cinerea* infection in WT, *ire1a/ire1b* and *bzip60* mutants. Plants were sprayed with ¼ PDB solution (Mock, white bars) or with ¼ PDB solution containing *B. cinerea* spores (black bars). For each experiments, four leaves of three plants were collected at 24 h (A) or 48 h (B) post-treatment. Transcript levels were quantified by RT-qPCR and normalized to the plant reference genes *AT4G26410* and *AT3G01150* transcript levels (Czechowski et al., 2005). The represented fold change is the mean of six independent experiments. Different letters represent groups which were significantly different from one another as determined by a one-way ANOVA followed by a multiple comparison with a Fisher’s Least Significant Difference (LSD) post-hoc test.

**Supplemental Figure S7.**
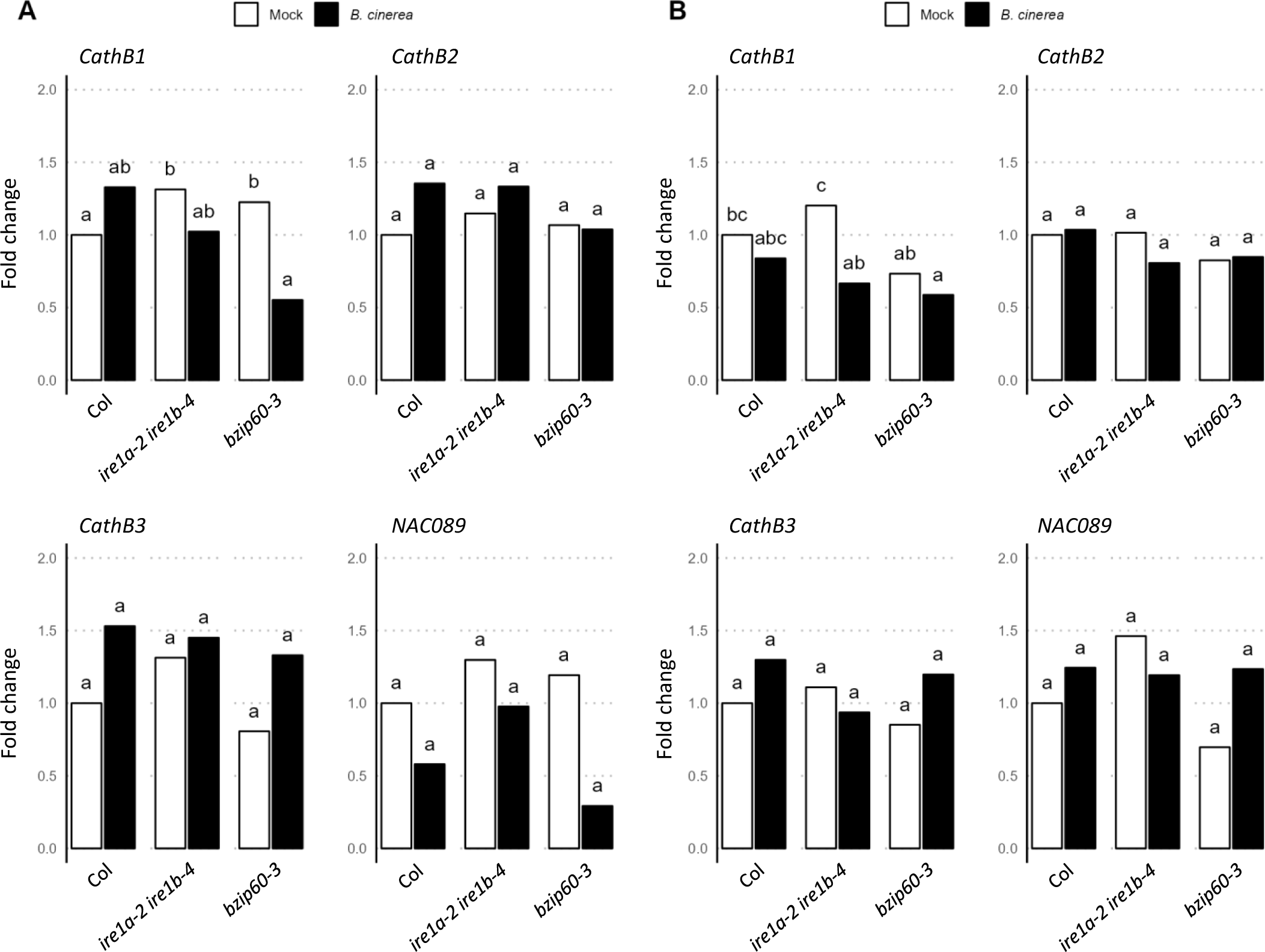
Accumulation of mRNA coding cathepsin and NAC089 transcription factor in response to *B. cinerea* infection in WT, *ire1a/ire1b* and *bzip60* mutants. Plants were sprayed with ¼ PDB solution (Mock, white bars) or with ¼ PDB solution containing *B. cinerea* spores (black bars). For each experiments, four leaves of three plants were collected at 24 h (A) or 48 h (B) post-treatment. Transcript levels were quantified by RT-qPCR and normalized to the plant reference genes *AT4G26410* and *AT3G01150* transcript levels (Czechowski et al., 2005). The represented fold change is the mean of six independent experiments. Different letters represent groups which were significantly different from one another as determined by a one-way ANOVA followed by a multiple comparison with a Fisher’s Least Significant Difference (LSD) post-hoc test.

**Supplemental Figure S8:**
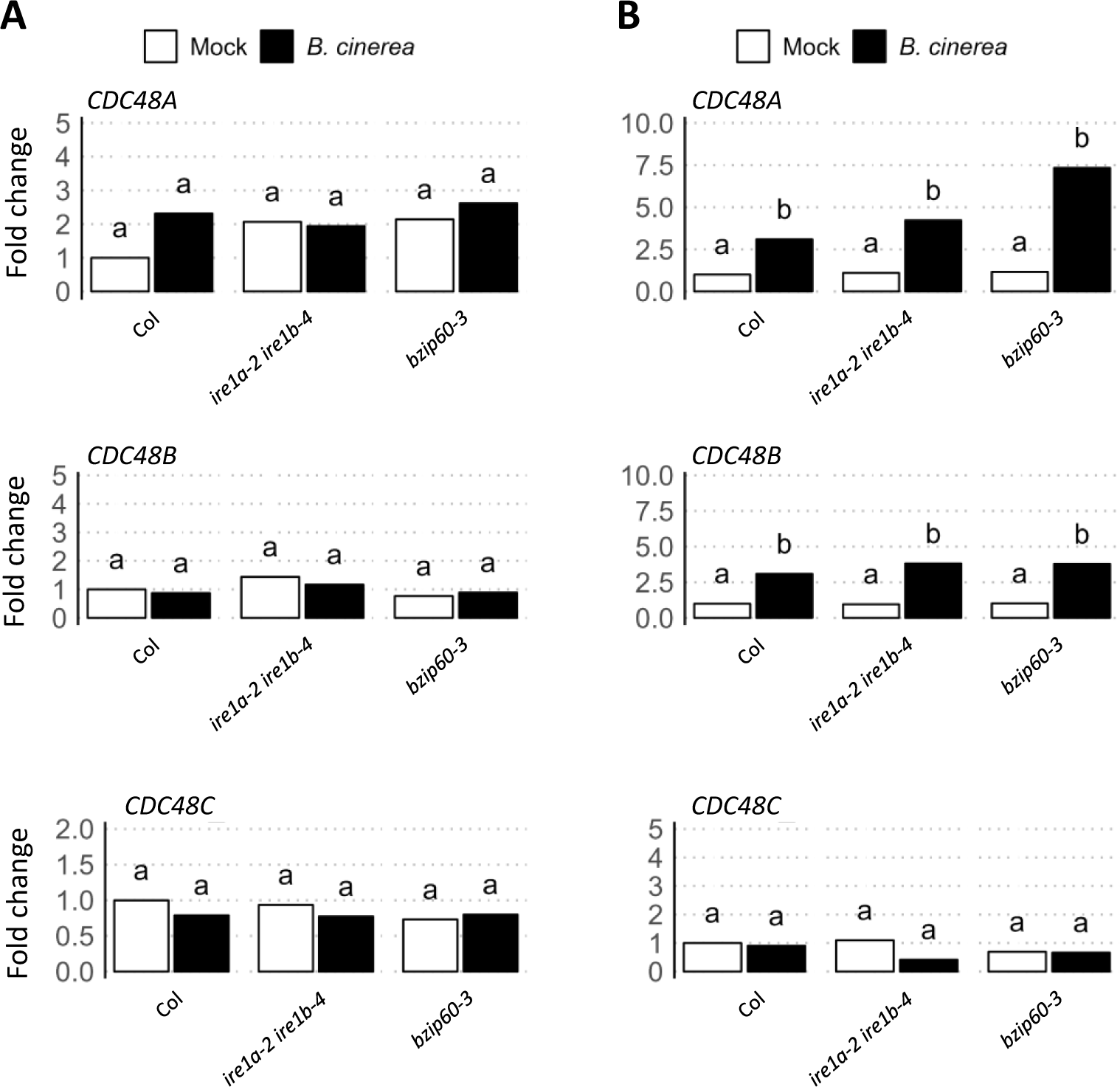
Accumulation of mRNA coding CDC48 in response to *B. cinerea* infection in WT, *ire1a/ire1b* and *bzip60* mutants. Plants were sprayed with ¼ PDB solution (Mock, white bars) or with ¼ PDB solution containing *B. cinerea* spores (black bars). For each experiments, four leaves of three plants were collected at 24 h (A) or 48 h (B) post-treatment. Transcript levels were quantified by RT-qPCR and normalized to the plant reference genes *AT4G26410* and *AT3G01150* transcript levels (Czechowski et al., 2005). The represented fold change is the mean of six independent experiments. Different letters represent groups which were significantly different from one another as determined by a one-way ANOVA followed by a multiple comparison with a Fisher’s Least Significant Difference (LSD) post-hoc test.

**Supplemental Figure S9:**
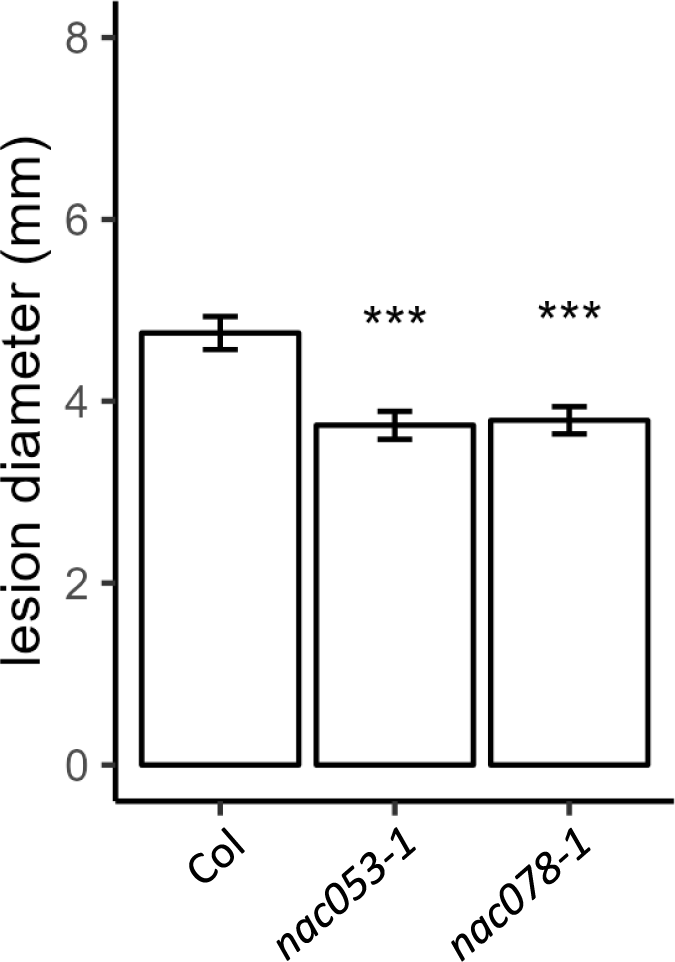
Mean lesion diameters observed on *nac053* and *nac078* mutants infected by *B. cinerea*. Lesion diameters observed on wild-type Col plants and mutants were measured 3 days after *B. cinerea* infection. Experiments were repeated four times. Seven plants per genotype were infected in each experiment. The stars indicate significant differences compared to the wild type which were identified using Kruskal-Wallis’ method followed by Dunnett’s post-hoc test(*** p<0.005).

**Supplemental Figure S10:**
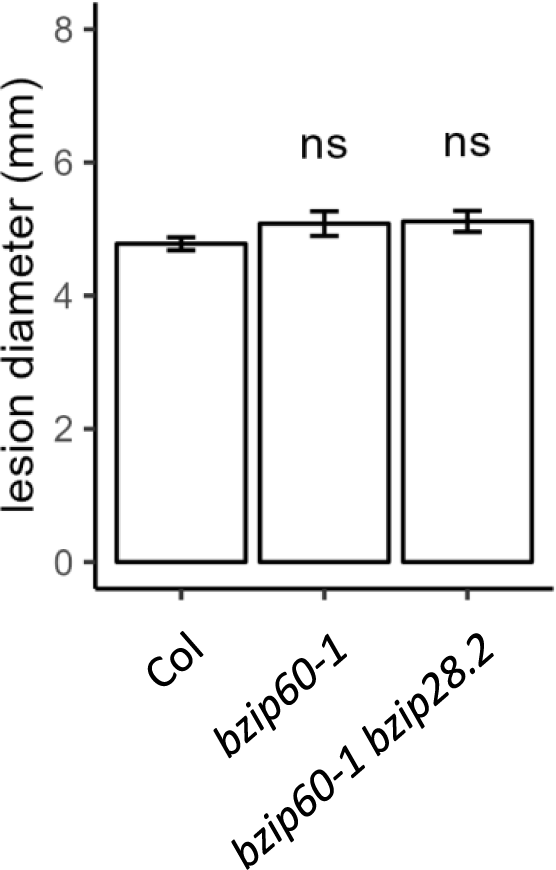
Mean lesion diameters observed on *bzip60.1-bzip28.2 double mutant* infected by *B. cinerea*. Lesion diameters observed on wild-type Col plants and mutants were measured 3 days after *B. cinerea* infection. Experiments were repeated six times. Seven plants per genotype were infected in each experiment. Significant differences compared to the wild type were identified using Kruskal-Wallis’ method followed by Dunnett’s post-hoc test (ns: not significant).

**Supplemental table S1:** List of mutants used in this study.

| mutant name | gene locus | mutant reference | references | origin |  |
| --- | --- | --- | --- | --- | --- |
| *Atbzip17-1* | At2g40950 | SALK_104326 | Liu et al., 2007a | Pr. Steven H Howell | https://doi.org/10.1111/j.1365-313X.2007.03195.x |
| *Atbzip28-2* | At3g10800 | SALK_132285C | Liu et al., 2007b | Pr. Steven H Howell | https://doi.org/10.1105/tpc.106.050021 |
| *Atbzip60-3* | At1g42990 | GABI_326A12 | Bao et al., 2012 | Pr. Steven H Howell | https://doi.org/10.1080/15548627.2018.1462426 |
| *Atire1a-2* | At2g17520 | SALK_018112 | Deng et al., 2011 | Pr. Steven H Howell | https://doi.org/10.1073/pnas.1102117108 |
| *Atire1b-4* | At5g24360 | SAIL_238_F07 | Deng et al., 2011 | Pr. Steven H Howell | https://doi.org/10.1073/pnas.1102117108 |
| *Atire1a-2/Atire1b-4* |  | SALK_018112 x SAIL_238_F07 | Moreno et al., 2012 | Pr. Steven H Howell | https://doi.org/10.1371/journal.pone.0031944 |
| *Aterdj3b-1* | At3g62600 | SALK_113364 | Nekrasov et al, 2009 | Pr. Cyril Zipfel | https://doi.org/10.1038/emboj.2009.262 |
| *Atsdf2.2* | At2g25110 | SALK_141321 | Nekrasov et al, 2009 | Pr. Cyril Zipfel | https://doi.org/10.1038/emboj.2009.262 |
| *Atcdc48a-4/muse8* | At3g09840 | EMS mutant | Copeland et al., 2016 | Pr. Xin Li | https://doi.org/10.1111/tpj.13251 |
| *Atcdc48b-3* | At3g53230 | Gabi_104F08 | our study | NASC |  |
| *Atcdc48b-4* | At3g53230 | Gabi_485G04 | our study | NASC |  |
| *Atcdc48c-2* | At5g03340 | SALK_102955C | our study | NASC |  |
| *Atcdc48c-3* | At5g03340 | SALK_123409 | our study | NASC |  |
| *Atnac53-1* | At3g10500 | SALK_009578C | Gladman et al., 2016 | NASC | https://doi.org/10.1105/tpc.15.01022 |
| *Atnac78-1* | At5g04410 | SALK_025098 | Gladman et al., 2016 | NASC | https://doi.org/10.1105/tpc.15.01022 |

NASC : Nottingham Arabidopsis Stock Centre (https://arabidopsis.info/)

**Supplemental table S2:**
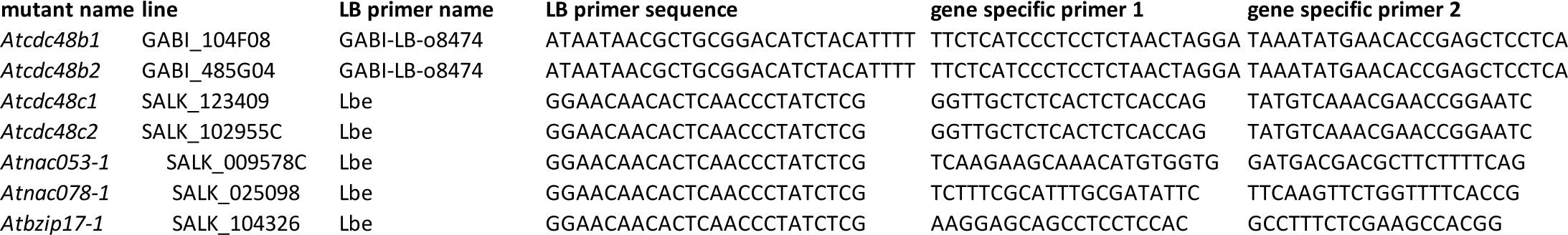
Primers used for mutant genotyping.

**Supplemental table S3:**
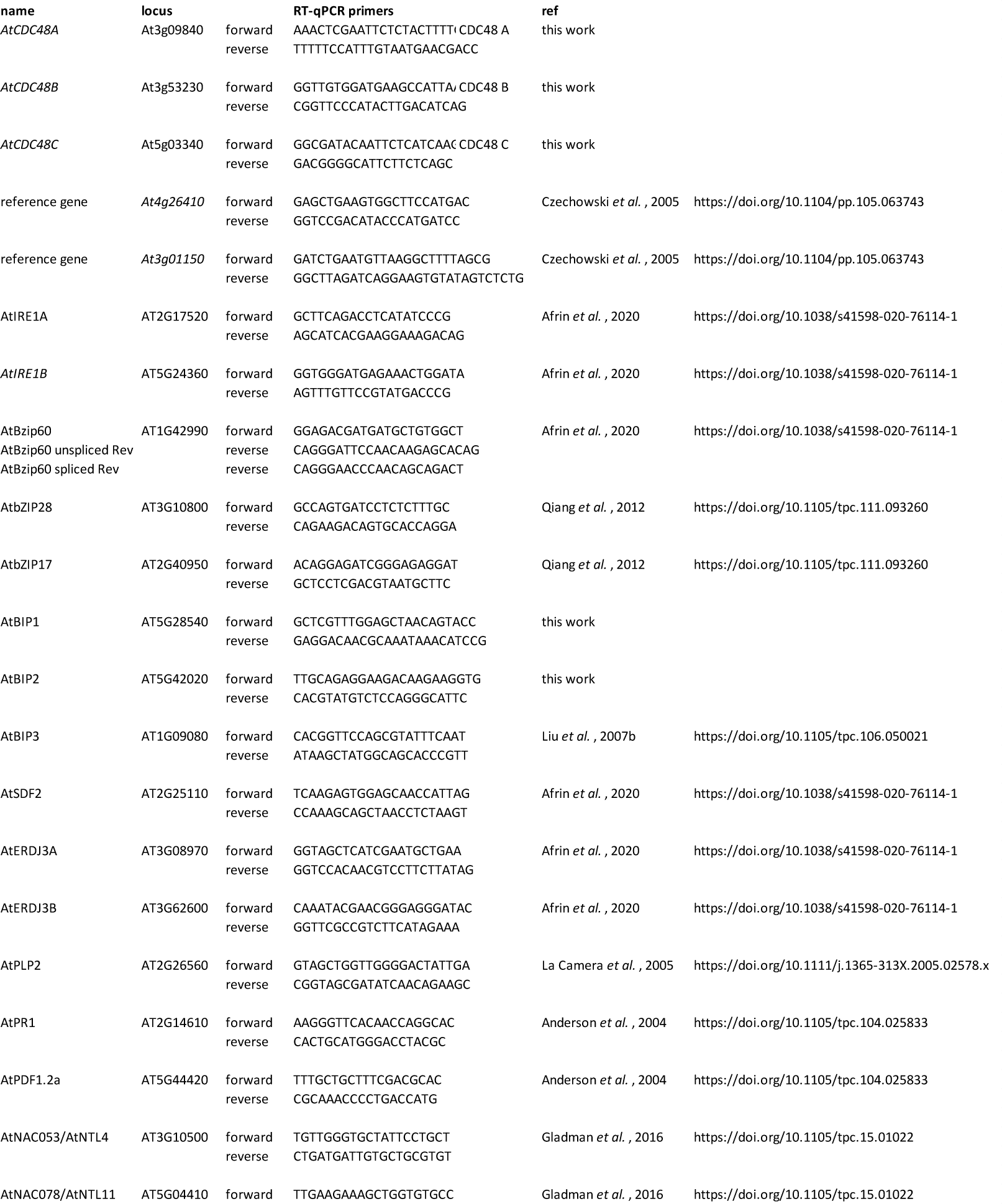
RT-qPCR primers used in this work.

## Notes

### Competing Interest Statement

The authors have declared no competing interest.

### Summary of Updates

- The text has been improved and mistakes corrected - The title has been updated. - “Supp Figure 4” has been changed to “Figure ”". - A new figure, “Figure 7,” has been added. - Former “Figure S5” has been split into “Figure S4” and “Figure S5”. - New figures “Figure S9” and “Figure S10” have been added.

